# Coordinated expansion of CD163⁺ monocytes and immature CD177⁺ neutrophils marks severe neurotoxicity after CD19 CAR T cell therapy

**DOI:** 10.64898/2026.07.07.737099

**Authors:** Tony Chour, Nikhita Poole, Hugh R. MacMillan, Katelyn Burleigh, David R. Glass, Emily C. Liang, Ryan Basom, Bobbie-Jo M. Webb-Robertson, Kelly Stratton, Derrik Gratz, Annalyssa N. Long, Anna E. Elz, Jennifer J. Huang, Alexandre V. Hirayama, Stanley R. Riddell, Jordan Gauthier, Heather H. Gustafson, Evan W. Newell, Sylvain Simon

**Author notes:** Corresponding Authors: Sylvain Simon, Translational Science and Therapeutics Division, Fred Hutchinson Cancer Center, 1201 Eastlake Ave E, Seattle, WA, 98109; Evan W. Newell, Vaccine and Infectious Disease Division, Fred Hutchinson Cancer Center, 1201 Eastlake Ave E, Seattle, WA 98109.

## Abstract

Immune effector cell-associated neurotoxicity syndrome (ICANS) is a major complication after CAR T cell therapy, but its underlying mechanisms remain poorly understood. We performed longitudinal immune profiling of paired whole blood and serum samples from patients with relapsed or refractory diffuse large B cell lymphoma (DLBCL) treated with CD19 CAR T cells. At peak neurotoxicity, high-dimensional mass cytometry and serum proteomics identified the expansion of CD163⁺ monocytes and immature CD10^low^CD101^low^ neutrophils correlated with elevated serum ST2 and IL-2RA concentrations. Integrative immune module analysis identified these features among the strongest predictors of ICANS severity. Independent single-cell transcriptomic profiling validated the emergence of immunoregulatory CD163⁺ monocytes and identified CD177 as a biomarker of ICANS-associated immature neutrophils. Together, these findings reveal a coordinated myeloid inflammatory network associated with ICANS and nominate candidate biomarkers and therapeutic targets for improving the safety of CAR T cell therapy.

**Significance:** We demonstrate that immunoregulatory CD163^+^ monocytes and immature, activated CD177^hi^CD10^low^CD101^low^ neutrophils emerge in patients with moderate to severe ICANS at peak toxicity following CD19 CAR T cell therapy. These findings identify an uncharacterized myeloid network potentially contributing towards ICANS pathogenesis.

## INTRODUCTION

Over the past decade, CAR T cell therapy has demonstrated its efficacy as a potent treatment option for relapsed B cell malignancies, including leukemia, lymphoma, and multiple myeloma^1^. Although CD19 CAR T-cell therapy has demonstrated remarkable efficacy in achieving remission, treatment-related toxicities commonly occur and can compromise clinical outcomes^2^. Approximately 70% of patients receiving CAR T-cell therapy develop cytokine release syndrome (CRS), with severe CRS reported in 8–42% of cases. Immune effector cell-associated neurotoxicity syndrome (ICANS) occurs in approximately 40% of patients and remains a major cause of treatment-related morbidity^3,4^. Although CRS and ICANS both correlate with elevated inflammatory serum proteins, including IL-6, IL-1β, MCP-1, and IFN-γ, ICANS remains a particularly challenging toxicity to treat because its pathobiology is less well defined and therapeutic options are limited. CRS is often responsive to IL-6 receptor blockade with tocilizumab and/or corticosteroids, whereas ICANS can be more prolonged and less consistently responsive to IL-6-directed therapies. As a result, treatment of ICANS commonly relies on corticosteroids and may incorporate additional immunomodulatory approaches, such as IL-1 receptor blockade with anakinra^5^. However, the cellular mechanisms that drive ICANS remain poorly defined. Therefore, we focused on defining immune cell populations associated with ICANS onset and severity to better understand the cellular mechanisms underlying neurotoxicity after CD19 CAR T cell therapy^6^. Furthermore, monocytes have been identified as a primary source of IL-6 during CRS and ICANS, highlighting the importance of integrating soluble mediator profiling with immune cell population analyses to define how coordinated changes in these compartments drive toxicity severity following CAR T-cell therapy^7^.

Prior studies have largely focused on characterizing activated and exhausted CAR T-cell phenotypes^8^ or quantifying CAR T-cell expansion and persistence following infusion as key correlates of therapeutic efficacy and safety^9^. While CAR T-cell research has traditionally focused on CAR T-cell expansion, persistence, and functional state, recent studies have implicated myeloid populations in treatment outcomes following CD19 CAR T-cell therapy^10^. Monocytes and neutrophils have emerged as important regulators of inflammatory responses in both CAR T-cell therapy and other hyperinflammatory conditions, including sepsis and COVID-19^11,12^. However, the specific myeloid populations associated with ICANS development remain poorly defined. Recent studies have drawn attention to the contributions of monocytes, macrophages and neutrophils to both clinical efficacy and treatment-related toxicities following CD19 CAR T cell therapy^13,14^. We hypothesize that neutrophils may also contribute towards inflammatory adverse events in CD19 CAR T cell therapy and seek to discover biomarkers characterizing these populations.

Historically, comprehensive characterization of granulocyte populations has been limited by their sensitivity to *ex vivo* manipulation and the technical challenges associated with profiling these cells using conventional single-cell approaches^15^. Whole-blood stabilization methods have enabled preservation of granulocyte phenotypes and support high-dimensional immunophenotyping directly from whole blood^16^. These approaches have facilitated more accurate assessment of granulocyte heterogeneity and longitudinal immune dynamics in clinical samples. In parallel, the development of fixed-cell compatible single-cell transcriptomic platforms, such as 10X Genomics FLEX APEX (GEM-X Flex v2), now enables transcriptional profiling of preserved granulocytes while minimizing artifacts introduced during sample processing. Together, these technologies provide complementary protein- and transcript-level measurements that allow comprehensive characterization of granulocyte states in longitudinal patient cohorts.

To investigate cellular and soluble mediators associated with toxicity following CD19 CAR T cell therapy, we collected longitudinal serum and whole-blood samples from patients with diffuse large B-cell lymphoma (DLBCL) treated with standard-of-care CD19-directed CAR T cells. Leveraging whole-blood-compatible mass cytometry and single-cell transcriptomic approaches, we characterized immune cell dynamics across treatment. In two independent cohorts, we identified the emergence of CD163⁺ monocytes and immature, activated CD177⁺CD10⁻ neutrophils in the blood of patients who developed moderate-to-severe immune effector cell–associated neurotoxicity syndrome (ICANS). These populations expanded near peak neurotoxicity and were associated with elevated inflammatory serum mediators, revealing a previously unrecognized myeloid network that may contribute to ICANS pathogenesis.

## RESULTS

### Whole blood immune composition of patients undergoing CD19 CAR T cell therapy is broadly heterogeneous relative to healthy donors

We longitudinally profiled whole blood samples from 33 patients with relapsed or refractory B-cell malignancies, including diffuse large B-cell lymphoma (DLBCL; n = 29), mantle cell lymphoma (MCL; n = 2), follicular lymphoma (FL; n = 1), and acute lymphoblastic leukemia (ALL; n = 1) **(Supplemental Table 1).** Our objective was to characterize immune correlates associated with adverse effects throughout the course of CD19 CAR T cell therapy. We focused on ICANS-associated correlations in this study as it is the less well-characterized adverse event in CD19 CAR T cell therapy. CRS and ICANS severity was graded based on the Lee 2019 consensus criteria and the Common Terminology Criteria for Adverse Events (CTCAE) v4.03 criteria^17^. Seventeen patients developed severe CRS, defined as grades 2-5, while sixteen developed moderate to severe ICANS, also defined as grade 2-5 **(Figure 1A)**. For patients who developed any grade CRS (n=30, grades 1-5), the median peak CRS toxicity was 5 days post-CAR T cell infusion (IQR 2-7). Among patients who developed any grade ICANS (n=23, grades 1-5), the median day for peak ICANS toxicity was 7 days post-CAR T cell infusion (IQR 4-10 days). Comprehensive blood count analysis demonstrated a global reduction in lymphocyte, mononuclear and neutrophil populations during the lymphodepletion (LD) and early post-infusion timeframe compared to baseline pre-LD levels (**Supplemental Figure 1A)**. Patients who developed moderate to severe ICANS had reduced peripheral basophil, eosinophil, monocyte and lymphocyte counts post CAR T cell infusion compared to patients that did not develop severe ICANS **(Figure 1A)**^18^. We observed elevated immature granulocyte counts within 20 days of CAR T-cell infusion in patients who subsequently developed moderate-to-severe ICANS compared with those who did not develop severe ICANS (**Figure 1A**; grade 0–1 ICANS, n=17; grade 2–5 ICANS, n=16).Monocyte counts remained reduced in patients who developed moderate to severe ICANS compared to those who did not at the time of peak neurotoxicity **(Supplemental Figure 1B-C)**.

**Figure 1.**
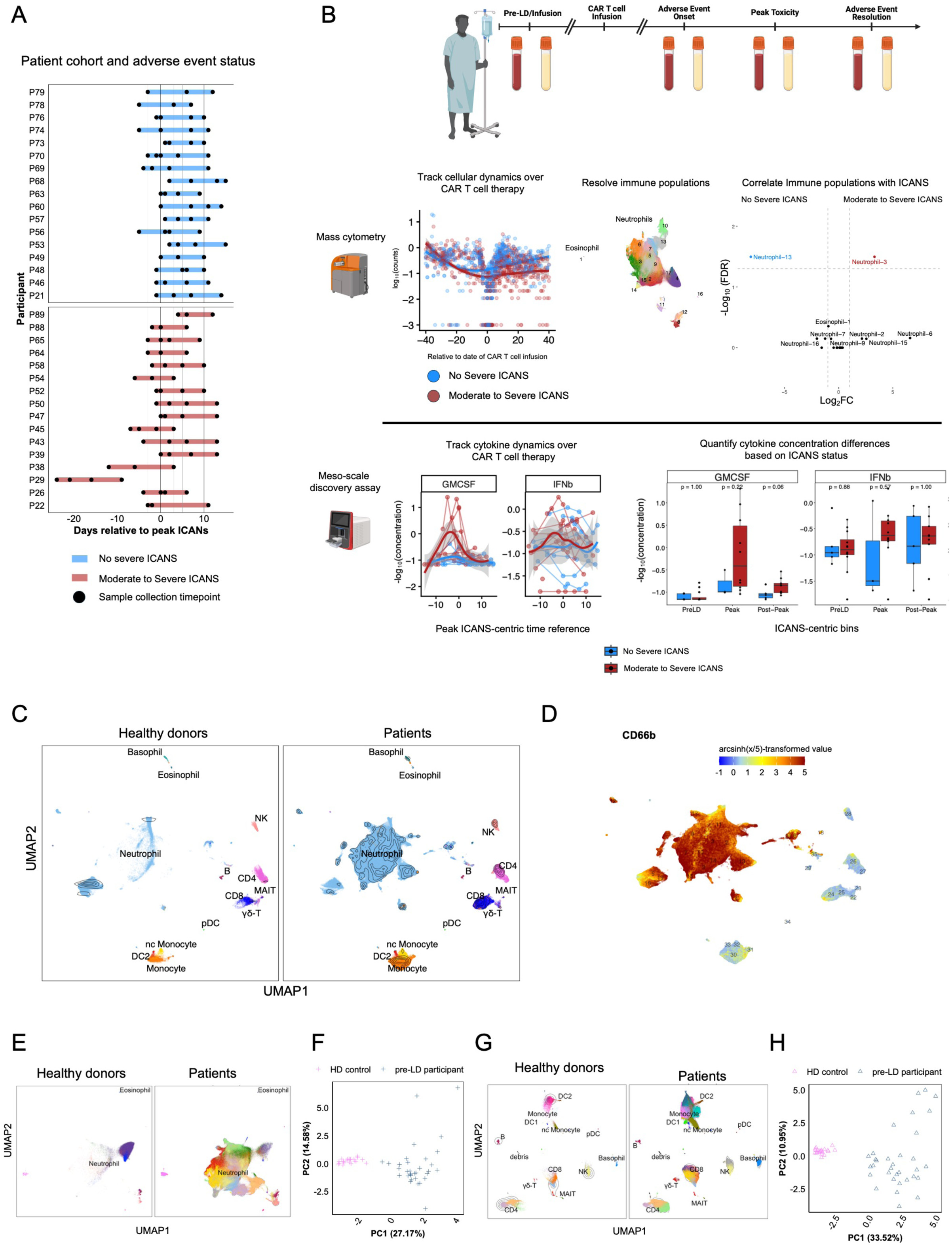
Study design and longitudinal immune profiling of patients undergoing CD19 CAR T-cell therapy. A. Swimmer plot depicting ICANS severity and sample collection timepoints for individual patients relative to peak ICANS onset (Day 0). For patients who did not develop ICANS, the cohort median peak ICANS onset day (Day 7 post-infusion) was used as the reference timepoint (n = 33). B. Schematic of longitudinal sample collection and experimental workflow. Fixed whole blood and serum samples were collected before lymphodepletion (PreLD) and at serial timepoints following CD19 CAR T-cell infusion. Whole blood samples were analyzed by mass cytometry (CyTOF) to profile innate and adaptive immune populations, while serum was used for soluble mediator quantification. C. Uniform manifold approximation and projection (UMAP) visualization of major immune populations identified across healthy donor controls and patient samples. D. CD66b expression across resolved immune populations highlighting granulocyte and non-granulocyte compartments. E. UMAP visualization of CD66b⁺ granulocyte populations from healthy donor controls and pre-lymphodepletion (PreLD) patient samples. F. Principal component analysis (PCA) of CD66b⁺ granulocyte populations demonstrating separation of healthy donor and PreLD patient samples. G. UMAP visualization of CD66b⁻ mononuclear populations from healthy donor controls and PreLD patient samples. H. Principal component analysis (PCA) of CD66b⁻ mononuclear populations demonstrating separation of healthy donor and PreLD patient samples.

Longitudinal profiling of 33 patients receiving CD19 CAR T-cell therapy revealed distinct cellular and soluble immune signatures associated with moderate-to-severe ICANS **(Figure 1B).**

We developed an integrated pipeline to quantify immune cell composition and serum soluble analyte concentrations at each sample collection timepoint. Whole blood contains diverse immune populations, including lymphocytes, monocytes, neutrophils, and other leukocytes. Using this approach, we (1) identified surface protein changes associated with moderate-to-severe ICANS, (2) resolved and characterized immune cell populations within the patient cohort, (3) defined biomarkers that distinguish these populations, and (4) tracked their longitudinal dynamics throughout CD19 CAR T-cell therapy to identify cellular changes associated with peak ICANS severity **(Figure 1B).**

Using a 40-marker mass cytometry panel encompassing lineage, activation, memory, proliferation, phosphorylation, and migration markers, we profiled peripheral leukocyte populations, including CD4⁺, CD8⁺, and γδ T cells, MAIT cells, B cells, monocytes, dendritic cells, natural killer cells, and granulocytes (neutrophils, eosinophils, and basophils), across patients receiving CD19 CAR T-cell therapy (n = 33) and healthy donors (n = 4) (**Figure 1C, Supplemental Table 2, Supplemental Figure 2).**

CD66b expression is restricted to neutrophil and eosinophil populations and was used for secondary, high-resolution clustering analysis of these populations **(Figure 1D)**. CD66b^+^ neutrophil and eosinophil clusters within healthy donors showed major compositional differences compared to patients **(Figure 1E)**. Principal component analysis demonstrated clear segregation between healthy donor and patient samples, with the first principal component (PC1) accounting for 27.17% of the total variance observed across granulocyte populations **(Figure 1F).** A homogeneous population of neutrophils dominated healthy donor profiles, while at baseline pre-LD collection timepoints, substantial population heterogeneity was observed within patient neutrophil samples. Similarly, among mononuclear CD66b^-^ cells, we observed a difference in the cellular composition populations between our healthy donor and patient Pre-LD samples (**Figure 1G)**. Principal component analysis again demonstrated separation between healthy donor and patient samples, with PC1 accounting for 33.52% of the total variance within the mononuclear compartment **(Figure 1H)**. We hypothesize that differences between healthy donor and Pre-LD patient samples within our neutrophil and mononuclear compartments may be driven by the underlying malignancy or history of chemotherapy prior to CAR T cell therapy. Additionally, disease-related inflammation and supportive care interventions such as glucocorticoid and G-CSF administration before CAR T-cell infusion, may contribute to the neutrophil and mononuclear cell heterogeneity observed at baseline.

### Distinct monocyte populations correlate with ICANS development

To characterize the diverse mononuclear cell populations within our patient cohort, we gated on CD66b⁻ cells and clustered cells into phenotypically distinct immune cell subsets **(Figure 2A).** Our mass cytometry panel enabled the resolution of 27 CD66b⁻ mononuclear clusters (2 potential debris/non-cell clusters) encompassing CD4 and CD8 T cells, NK cells, B cells, monocytes, and dendritic cells subsets **(Figure 2B)**.

**Figure 2.**
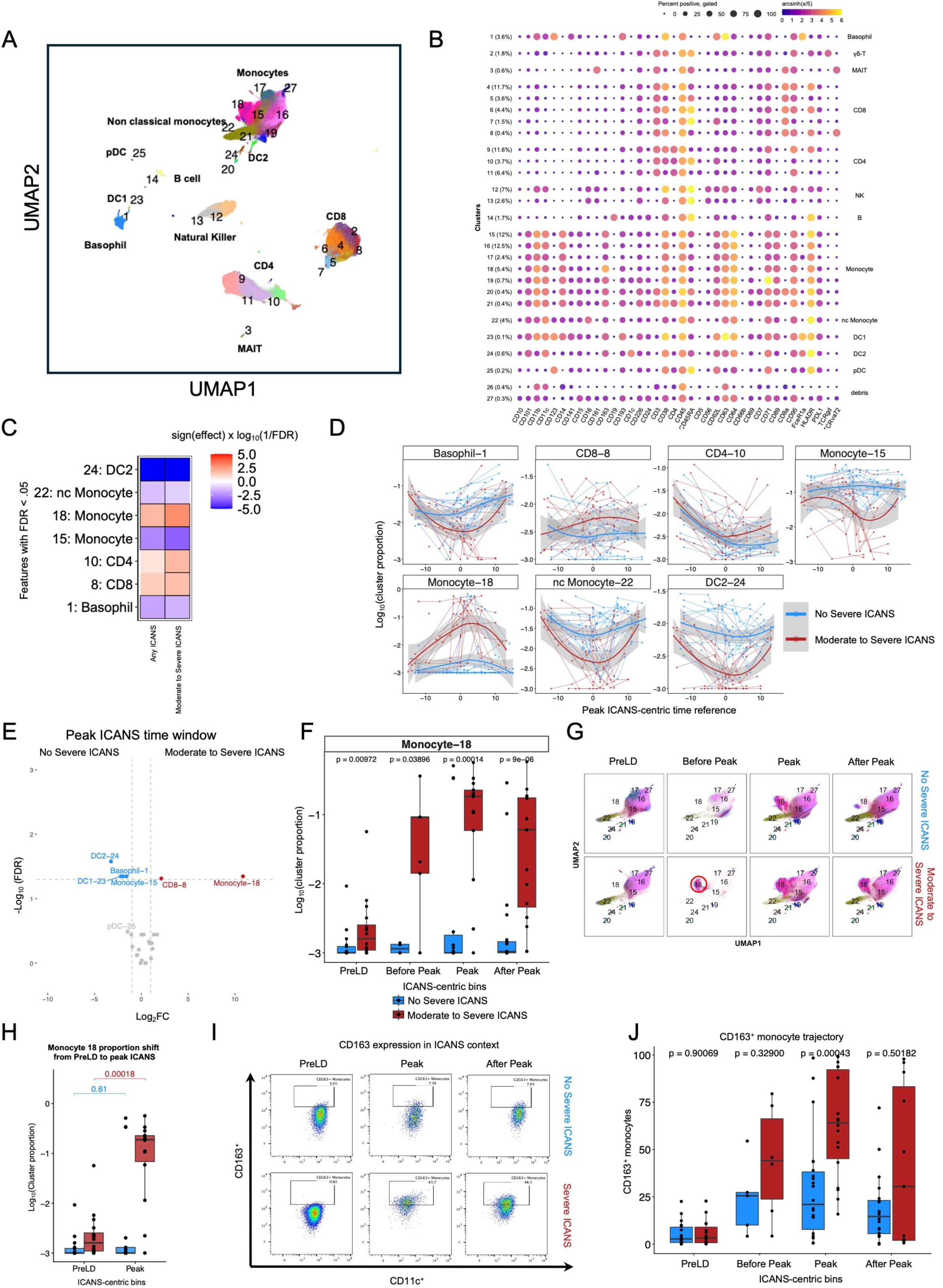
Longitudinal mononuclear profiling identifies CD163⁺ monocytes associated with moderate-to-severe ICANS. A. Uniform manifold approximation and projection (UMAP) visualization of CD66b⁻ mononuclear populations identified across all patient and healthy donor samples. B. Bubble plot showing median surface marker expression used to annotate CD66b⁻ mononuclear populations. C. Associations between mononuclear populations and CRS or ICANS severity identified using generalized estimating equations (GEE). Outlined populations represent statistically significant associations after false discovery rate (FDR) correction (FDR < 0.05). D. Longitudinal abundance of mononuclear populations expressed as a proportion of CD66b⁻ cells relative to peak ICANS onset (Day 0). For patients who did not develop ICANS, the cohort median peak ICANS onset day (Day 7) was used as the reference timepoint. Lines connect serial samples collected from the same patient. E. Differentially abundant mononuclear populations at peak ICANS comparing patients with non-severe ICANS (grade 0–1) and moderate-to-severe ICANS (grade 2–5). Colored populations indicate statistical significance following FDR correction (FDR < 0.05; Wilcoxon rank-sum test). F. Temporal changes in Monocyte-18 abundance across ICANS-centric time windows in patients with non-severe and moderate-to-severe ICANS. G. Representative UMAPs showing mononuclear population composition across ICANS-centric time windows from a patient with non-severe ICANS (top) and a patient with moderate-to-severe ICANS (bottom). H. Change in Monocyte-18 abundance from baseline (PreLD) to peak ICANS in patients with non-severe and moderate-to-severe ICANS. I. Representative manual gating strategy identifying CD163⁺CD11c⁺ monocytes within the CD14⁺ monocyte compartment. J. Frequency of manually gated CD163⁺ monocytes across ICANS-centric time windows in patients with non-severe and moderate-to-severe ICANS.

We applied generalized estimating equations (GEE) to model the longitudinal abundances and look for differences associated with ICANS gradations and moderate to severe ICANS development within mononuclear clusters after CD19 CAR T cell infusion **(Figure 2C)**. Among the 27 mononuclear clusters identified, seven were associated (FDR < 0.05) with graded ICANS severity or moderate to severe ICANS development. These seven mononuclear populations included basophils (basophil-1), dendritic cells (DC2-24), non-classical CD16^+^ monocytes (nc Monocyte-22), CD14^+^ monocytes (Monocyte 15 and Monocyte 18), CD4 T cells (CD4-10) and CD8 T cells (CD8-8).

While GEE captured broad ICANS associations throughout the entire course of CAR T cell therapy, we were interested in temporally resolved differences surrounding peak neurotoxicity. To characterize biomarkers at peak neurotoxicity, we aligned sample timepoints based on each patient’s estimated day of peak ICANS (**Figure 2D, Supplemental Figure 3**). Specifically, for patients with ICANS, we discretized time into four bins: 1) baseline pre-lymphodepletion—cells derived from samples collected prior to lymphodepletion and CAR T cell infusion; 2) Before Peak ICANS—spanning infusion date to three days preceding peak ICANS onset 3) Peak ICANS, defined as samples collected within ±3 days of the day of peak ICANS severity; and 4) After Peak ICANS—defined as >3 days after peak onset. For patients without severe ICANS, day 7 post-infusion (the median day of peak ICANS onset in our patients) was used as a reference point. In each designated timeframe, we identified mononuclear populations enriched in patients who developed moderate to severe ICANS **(Supplemental Figure 4A-B).**

Within this framework, we observed that moderate to severe ICANS patients exhibited elevated proportions of a CD163^+^ monocyte population (Monocyte-18) during peak toxicity relative to patients who do not develop severe ICANS **(Figure 2E).** Notably, Monocyte-18 proportions rose throughout the course of CD19 CAR T cell therapy and remained elevated in moderate to severe ICANS patients post-peak ICANS development relative to patients who do not experience severe ICANS **(Figure 2F-G)**. Notably, Monocyte-18 abundance positively correlated with cumulative dexamethasone exposure administered from CAR T-cell infusion through peak ICANS, but did not correlate with cumulative methylprednisolone or filgrastim exposure **(Supplemental Figure 5A–F).**

Among patients that developed moderate to severe ICANS, Monocyte-18 abundance significantly increased within the blood mononuclear from the baseline Pre-LD levels to Peak ICANS levels compared to patients without severe ICANS **(Figure 2H)** (Log10FC = 2.14, p =.00018 for our moderate to severe ICANS patients compared to Log10FC = 0.587,and p = .61 for non-severe ICANS patients, n= 14 and n= 17 respectively). Manual gating to quantify CD163^+^ populations within our CD14^+^ monocytes verified that CD163^+^ monocytes were elevated in patients who develop moderate to severe ICANS at the time of peak ICANS onset **(Figure 2I).** In addition to this peak-associated increase, CD163⁺ monocytes frequencies exhibited a progressive upward trajectory preceding peak ICANS onset in moderate to severe cases **(Figure 2J)**.

Beyond ICANS, this CD163^+^ monocyte population also positively correlated with severe CRS within the timeframe of peak CRS relative to other mononuclear populations (**Supplemental Figure 6A-B**). Together, these findings support CD163^+^ monocytes as a strong correlate of CRS and ICANS in CD19 CAR T cell therapy.

In contrast to the CD163^+^ monocyte population associated with moderate to severe ICANS, we observed a consistent negative correlation between moderate to severe ICANS and the abundance of classical monocyte, non-classical monocytes, and monocyte-derived dendritic cells during the peak and post-peak ICANS timeframes (Monocyte-15, nc Monocyte-22, and DC2-24 respectively) **(Figure 2D)**.

Together, these findings indicate that moderate-to-severe ICANS is characterized by distinct remodeling of the monocyte compartment, marked by the expansion of CD163⁺ monocytes and a concomitant reduction in classical monocyte and monocyte-derived dendritic cell populations.

### Immature neutrophils are positively correlated with the peak development of ICANS

In addition to mononuclear cell correlates of adverse events following CAR T-cell therapy, several neutrophil-associated serum proteins have been reported to increase during CRS, suggesting that neutrophil activation or dysregulation may also contribute to the development of treatment-related toxicities^19^. However, it remains unclear whether changes in neutrophil composition are associated with ICANS severity, highlighting the need to characterize neutrophil population dynamics during CAR T-cell therapy. To address this question, we profiled CD66b^+^ cells with mass cytometry to resolve distinct populations and identify neutrophils and eosinophils subsets associated with ICANS onset and severity.

Our clustering analysis resolved 17 distinct CD66b^+^ populations, including one eosinophil population and 16 neutrophil subsets **(Figure 3A-B)**. Neutrophil subpopulations exhibit continuous, gradient-like marker expression in contrast to discrete lineage-defined expression observed in mononuclear populations **((Figure 3C, Supplemental Figure 7)**.

**Figure 3.**
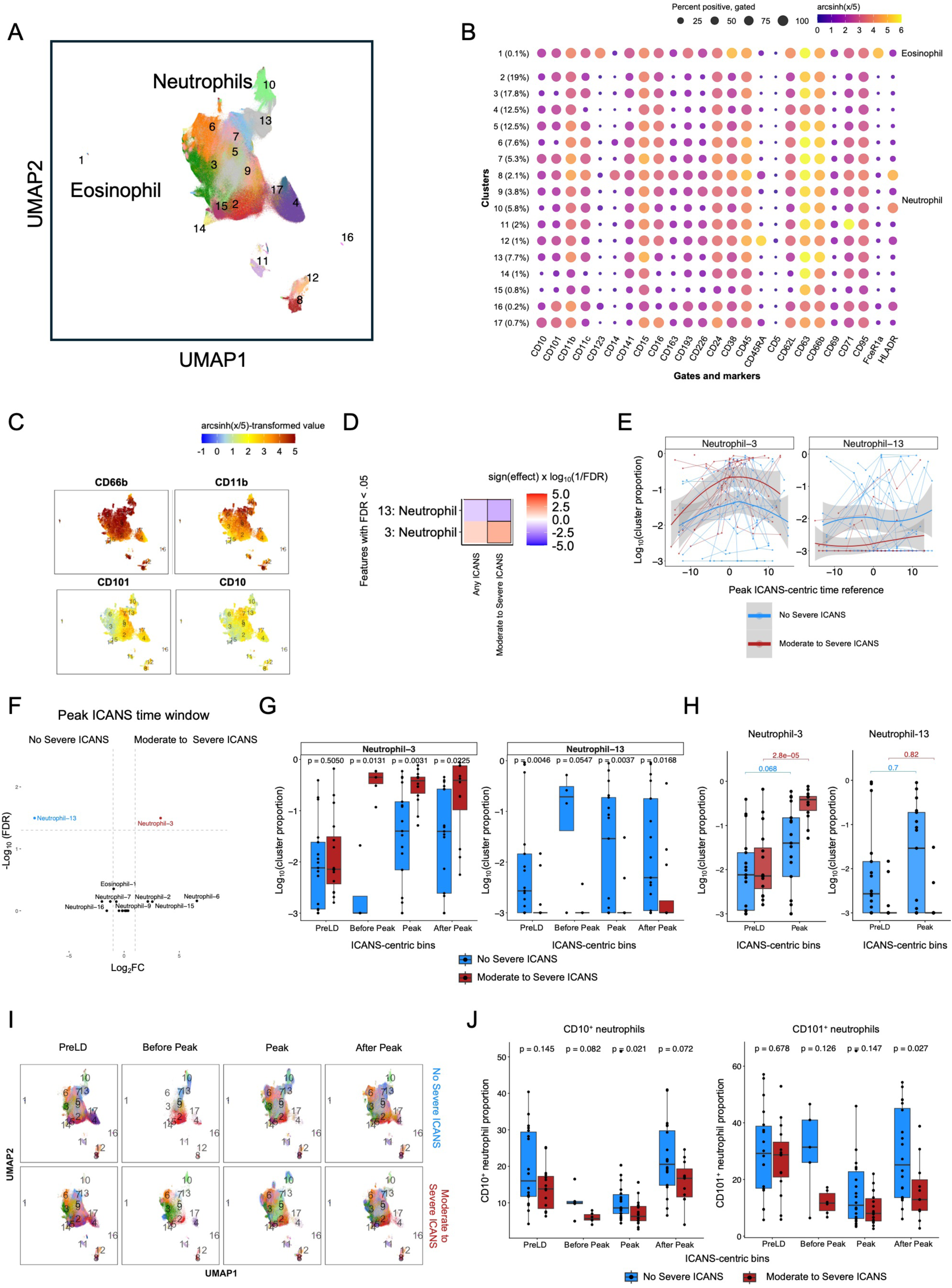
Longitudinal granulocyte profiling reveals CD10- CD101^-^ neutrophils assocaited with moderate to severe ICANS. A. Uniform manifold approximation and projection (UMAP) visualization of CD66b⁺ granulocyte populations identified across all patient and healthy donor samples. B. Bubble plot showing median surface marker expression used to annotate CD66b⁺ granulocyte populations. C. Feature plots displaying expression of selected surface proteins across granulocyte populations.Z D. Associations between granulocyte populations and ICANS severity identified using generalized estimating equations (GEE). Outlined populations represent statistically significant associations following false discovery rate (FDR) correction (FDR < 0.05). E. Longitudinal abundance of selected granulocyte populations expressed as a proportion of CD66b⁺ cells relative to peak ICANS onset (Day 0). Patients with moderate-to-severe ICANS (grade 2–5) are shown in red and patients with non-severe ICANS (grade 0–1) in blue. For patients who did not develop ICANS, the cohort median peak ICANS onset day (Day 7) was used as the reference timepoint. Lines connect serial samples collected from the same patient. F. Differentially abundant granulocyte populations at peak ICANS comparing patients with non-severe ICANS and moderate-to-severe ICANS. Colored populations indicate statistical significance following FDR correction (FDR < 0.05; Wilcoxon rank-sum test). G. Abundance of Neutrophil-3 and Neutrophil-13 across ICANS-centric time windows in patients with non-severe and moderate-to-severe ICANS. H. Change in Neutrophil-3 and Neutrophil-13 abundance from baseline (PreLD) to peak ICANS in patients with non-severe and moderate-to-severe ICANS. I. Representative UMAPs showing granulocyte population composition across ICANS-centric time windows in patients with non-severe ICANS (top) and moderate-to-severe ICANS (bottom). J. Quantification of CD10^low^ and CD101^low^ neutrophil populations within the CD66b⁺ compartment in patients with non-severe ICANS and moderate-to-severe ICANS, measured by mean fluorescence intensity (MFI).

We used GEE to test for granulocyte clusters associated with ICANS severity over all post-infusion timepoints, centered at peak ICANS **(Supplemental Figure 8).** We tested ICANS severity grades, as well as dichotomized (grade >1 or not) ICANS severity. Among the 17 CD66b^+^ populations, two neutrophil clusters were significantly associated with ICANS outcomes (FDR < 0.05) (Neutrophil-3 and Neutrophil-13) **(Figure 3D)**. Neutrophil-3 was characterized by low expression of both CD10 and CD101 relative to the other neutrophil clusters. Notably, Neutrophil-3 exhibited a positive association with moderate to severe ICANS at the time of peak toxicity, whereas Neutrophils-13 demonstrated negative association. **(Figure 3E, Supplemental Figure 9A-B)**. We also observed the emergence of an HLA-DR^+^ neutrophil population (Neutrophil-10) that was elevated during the peak ICANS window in patients regardless of ICANS status **(Supplemental Figure 8).**

Within each discrete time window, we observed the most pronounced difference in Neutrophil 3 proportions at peak ICANS, with markedly higher levels in patients who developed moderate to severe ICANS compared with those who did not **(Figure 3F-G).** Although Neutrophil 3 was the only neutrophil cluster to show a statistically significant correlation with moderate to severe ICANS at peak neurotoxicity, three additional clusters (Neutrophils-2, Neutrophils-6, and Neutrophils-15) trended toward enrichment with moderate to severe ICANS onset during this same period **(Figure 3F).** These moderate to severe ICANS trending clusters, in addition to Neutrophil-3 cluster, all exhibited low CD10 and to a lesser extent, low CD101 expression levels **(Figure 3C)**. CD10 and CD101 have been described as maturity markers in other inflammatory contexts, such as sepsis and COVID-19^20^ suggesting that the expansion of CD10^low^/CD101^low^ neutrophils may reflect the accumulation of immature inflammatory neutrophils in patients who develop severe ICANS. Notably, we did not observe a correlation between Neutrophil-3 abundance at peak ICANS and treatments administered (dexamethasone, methylprednisolone, or G-CSF (filgrastim) at the start of CAR T cell infusion through peak ICANS **(Supplemental Figure 10A-F).** Among patients who developed severe ICANS, Neutrophil-3 abundance significantly increased from baseline (Pre-LD) to peak ICANS onset **(Figure 3H-I),** with a log10 fold-change of 1.49 compared to 0.85 in patients without severe ICANS (n = 14 and n = 17, respectively). In contrast, Neutrophil-13 abundance, which was inversely associated with moderate to severe ICANS, did not significantly increase between Pre-LD and peak ICANS in severe cases. **(Figure 3H-I)**.

In parallel with unsupervised clustering analyses, we quantified CD10 mean fluorescence intensity (MFI) within manually gated total neutrophils across longitudinal time bins to validate changes in neutrophil maturation state at the population level. At peak ICANS, we observed significant reduced CD10 expression, consistent with enrichment of immature neutrophils in patients who developed moderate to severe ICANS at peak toxicity. Although neutrophil CD101 expression was lower at peak toxicity in patients who developed moderate-to-severe ICANS than in those without severe ICANS, the difference was not statistically significant. In both groups, CD10 and CD101 expression progressively declined during CAR T-cell therapy and reached their lowest levels around the time of peak ICANS. (**Figure 3J)**.

Together, these results demonstrate that ICANS is associated with distinct alterations in the neutrophil compartment, characterized by the expansion of CD10^low^CD101^low^ neutrophil populations and reduced expression of neutrophil maturation markers. The enrichment of these immature neutrophil subsets at peak neurotoxicity suggests that neutrophil maturation state may serve as an important correlate of ICANS severity following CD19 CAR T-cell therapy.

### Patients who develop severe ICANS have distinct cytokine signatures at peak ICANS

In parallel with cellular profiling, we characterized serum proteomic signatures associated with moderate to severe ICANS development. We designed ELISA and multiplex MesoScale Discovery (MSD) panels to measure a broad range of inflammatory cytokines pertinent to CD19 CAR T cell therapy, including GM-CSF, IL-6, interferon-gamma (IFN-γ), members of the IL-1 cytokine family such as IL-18, and multiple chemokines of interest (**Supplemental Table 3)**. We quantified serum cytokine levels in 9 patients who did not develop severe ICANS and 14 patients who developed moderate to severe ICANS.

We prioritized correlating soluble mediators with ICANS rather than CRS as soluble mediators associated with ICANS are not as well characterized. In addition, interpretation of CRS-associated cytokine patterns may have been confounded by tocilizumab administration in a substantial proportion of patients (16/33). GEE analysis identified and validated ST2 and IL-6 as longitudinal correlates of ICANS and further revealed Eotaxin-2 and Eotaxin-3 as novel soluble factors associated with ICANS across the course of CAR T-cell therapy **(Figure 4A)**^21,22^. To further define temporal cytokine dynamics surrounding neurotoxicity onset, we aligned patient samples relative to estimated peak ICANS dates, or day 7 post-infusion for patients without neurotoxicity. Cytokines associated with peak ICANS were then assessed longitudinally according to ICANS severity status **(Figure 4B, Supplemental Figure 11A-B).** At peak ICANS, serum protein concentrations were quantified from 10 patients without severe ICANS and 6 patients with severe ICANS. Eotaxin-3, ST2, and IL2RA were significantly elevated in patients with moderate to severe ICANS **(Figure 4C**). Discrete-time analyses further demonstrated that Eotaxin-3, IL2RA, and ST2 levels increased from pre-lymphodepletion baseline to peak ICANS in patients who developed moderate to severe ICANS and were significantly higher at the peak ICANS compared to patients who did not develop severe ICANS **(Figure 4D).**

**Figure 4.**
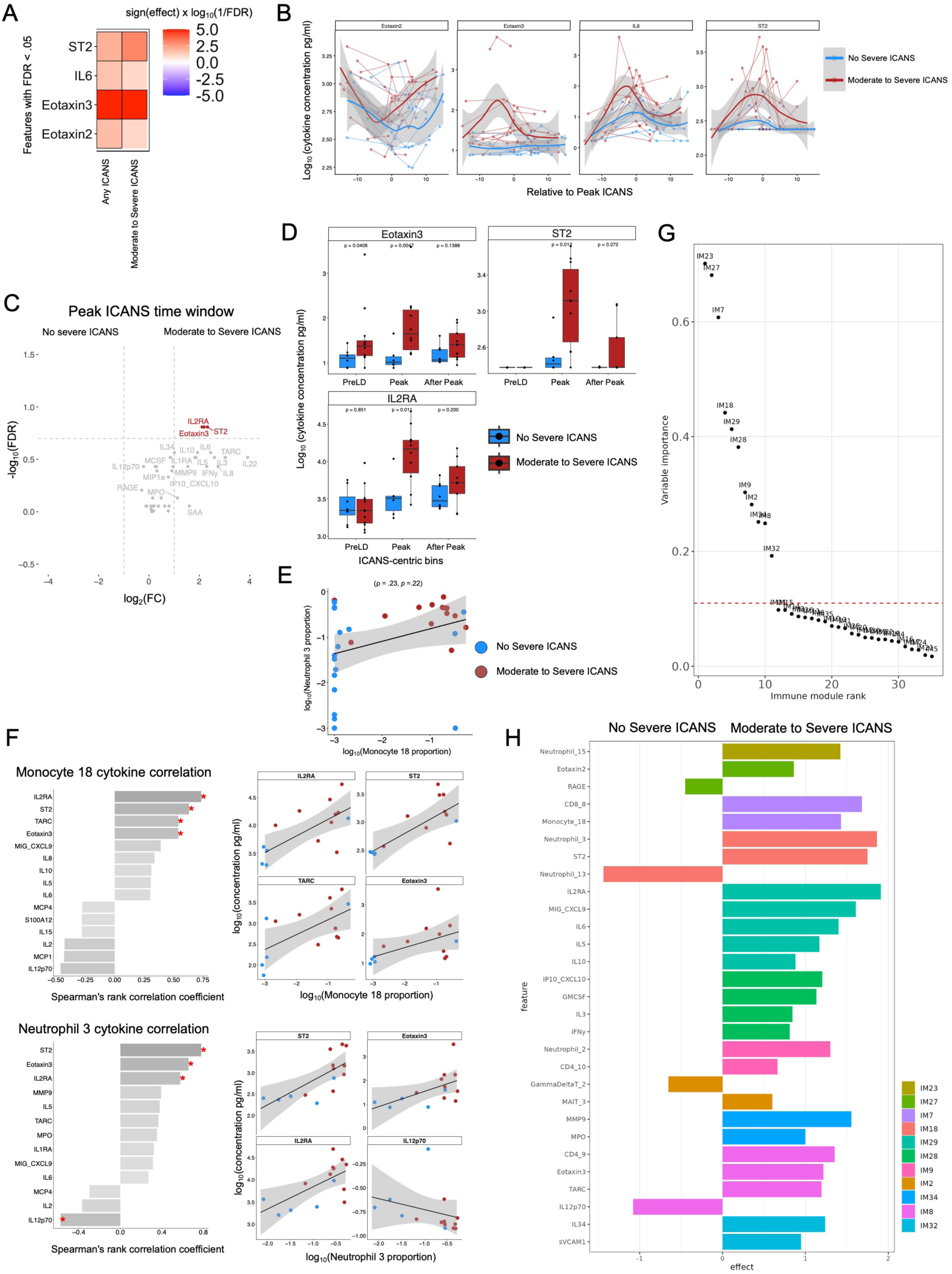
Soluble mediator profiling identifies ST2 and IL2RA as biomarkers of moderate-to-severe ICANS and reveals coordinated immune signatures associated with toxicity. A. Associations between circulating soluble mediators and ICANS severity identified using generalized estimating equations (GEE). Outlined analytes represent statistically significant associations following false discovery rate (FDR) correction (FDR < 0.05). B. Longitudinal soluble mediator concentrations relative to peak ICANS onset (Day 0). For patients who did not develop ICANS, the cohort median peak ICANS onset day (Day 7) was used as the reference timepoint. Lines connect serial samples collected from the same patient. C. Differentially abundant soluble mediators at peak ICANS comparing patients with non-severe ICANS (grade 0–1) and moderate-to-severe ICANS (grade 2–5). Colored analytes indicate statistical significance following FDR correction (FDR < 0.05; Wilcoxon rank-sum test). D. Longitudinal changes in selected soluble mediator concentrations throughout CAR T-cell therapy. E. Abundance of Monocyte-18 and Neutrophil-3 populations measured at peak ICANS. F. Associations between soluble mediator concentrations and Monocyte-18 or Neutrophil-3 abundance at peak ICANS assessed by linear regression. G. Predictive performance of individual immune features for identifying patients who developed moderate-to-severe ICANS. H. Effect sizes of features comprising the highest-performing immune modules predictive of moderate-to-severe ICANS.

In summary, these findings demonstrate that patients who develop moderate-to-severe ICANS exhibit a distinct systemic inflammatory signature characterized by elevated Eotaxin-3, ST2, and IL2RA at peak neurotoxicity.

### Immune modules elucidate biomarkers predictive of severe ICANS development in CD19 CAR T cell therapy

For a comprehensive systems-level perspective on the cellular and molecular dynamics driving moderate to severe ICANS development in CD19 CAR T cell therapy, we integrated our clustering analysis with our cytokine analysis. We examined correlations between CD66b⁺ neutrophil clusters, CD66b⁻ mononuclear clusters, and circulating cytokines at peak ICANS **(Supplemental Figure 12)**. We observed that patients who develop moderate to severe ICANS tend to exhibit higher proportions of both Monocyte-18 and Neutrophil-3, suggesting a coordinated expansion that may be associated with moderate to severe ICANS **(Figure 4E).**

To identify inflammatory mediators associated with CD163⁺ monocytes and immature neutrophils at peak ICANS, we performed Spearman correlation analyses of Monocyte-18 and Neutrophil-3 cluster frequencies and serum protein concentrations **(Supplemental Figure 13A-B).** Monocyte-18 was most positively associated with IL-2RA (ρ = 0.735, *p* = 0.002) and ST2 (ρ = 0.79, *p* = 0.005). Similarly, Neutrophil 3 was also most positively correlated with peripheral ST2 (ρ = 0.779, *p* = 0.001) and IL-2RA (ρ = 0.578, *p* = 0.03, and is inversely associated with peripheral IL-12p70 (ρ = −0.576, *p* = 0.03) **(Figure 4F).** The three immature neutrophil populations, Neutrophil-2, Neutrophil-6, and Neutrophil-15, displayed distinct cytokine correlation profiles, underscoring functional heterogeneity within this compartment **(Supplemental Figure 12).**

To interpret these coordinated cellular and cytokine dynamics, we applied an immune modules computational framework to integrate our multi-omic dataset and assess systems-level interactions during peak ICANS^23^. This approach clusters correlated serum cytokines and cellular phenotypes at each time point into immune modules, summarizes their variance using PCA, and uses the summarized immune modules as inputs into a random forest classifier predicting patient ICANS status.

From the peak ICANS time bin, we resolved 40 immune modules across all features within the 16 patients that had complete MSD serum and CyTOF cellular datasets and highlighted the most predictive immune modules for downstream analysis **(Figure 4G).** The number of immune modules resolved was based on maximizing the silhouette score. From ranked predictive modules, Neutrophil-15 emerged as the top predictor of moderate to severe ICANS onset. Neutrophil-15 was characterized as an immature CD10^low^CD101^low^ neutrophil population from our mass cytometry panel. We also observed our two cellular populations of interest, Monocyte-18 and Neutrophil-3 as top predictors of moderate to severe ICANS onset. ST2 and Neutrophil-3 were grouped into the same immune module, implying a strong correlation within these features strongly associated with moderate to severe ICANS during the critical ICANS window **(Figure 4H).**

Overall, the immune modules approach revealed stronger soluble protein-protein correlations and cellular-cellular population correlations than protein-cell correlations at the peak ICANS timepoint. We observe GM-CSF grouped together with IL-3, consistent with their shared cytokine family membership and overlapping signaling pathways. Taken together, these findings suggest a dynamic and interconnected cellular and proteomic network that shapes toxicity outcome following CD19 CAR T cell therapy.

### CD163^+^ monocytes are characterized by anti-inflammatory and wound healing gene signatures

To validate our mass cytometry findings, we performed single-cell transcriptomic profiling on a second cohort of 12 patients with relapsed or refractory diffuse large B-cell lymphoma (DLBCL) mantle cell lymphoma (MCL) or follicular lymphoma (FH) treated with the CD19 CAR T-cell product lisocabtagene maraleucel (Supplemental Table 1). Three patients developed moderate-to-severe ICANS (grades 2–5), while nine did not. CRS occurred in nine patients, including one patient with grade 2 CRS. Whole blood was collected longitudinally at Pre-LD and on Days 0, 1, 3, 7, 10, and 14 following CAR T-cell infusion. Samples were fixed with 4% paraformaldehyde and processed with the 10X FLEX APEX platform for single-cell transcriptomic analysis.

Using the FLEX APEX targeted probe set together with a custom Woodchuck Hepatitis Virus Post-Transcriptional Regulatory Element (WPRE) probe for detection of CAR T cells, 43 cellular clusters were resolved, including CD4⁺ and CD8⁺ T cells, CAR T cells, B cells, CD14⁺ and CD16⁺ monocytes, granulocytes, and platelets (**Figure 5A-B**). In addition to lineage markers, we broadly profile cytokine secretion capacity across our immune populations resolved **(Supplemental Figure 14).** To further characterize monocyte heterogeneity, we selected monocytic clusters based on *CD14, CCR2, FCGR3A* and *CSF1R* expression and performed secondary clustering to achieve higher-resolution identification of monocyte subpopulations **(Figure 5 B-C).**

**Figure 5.**
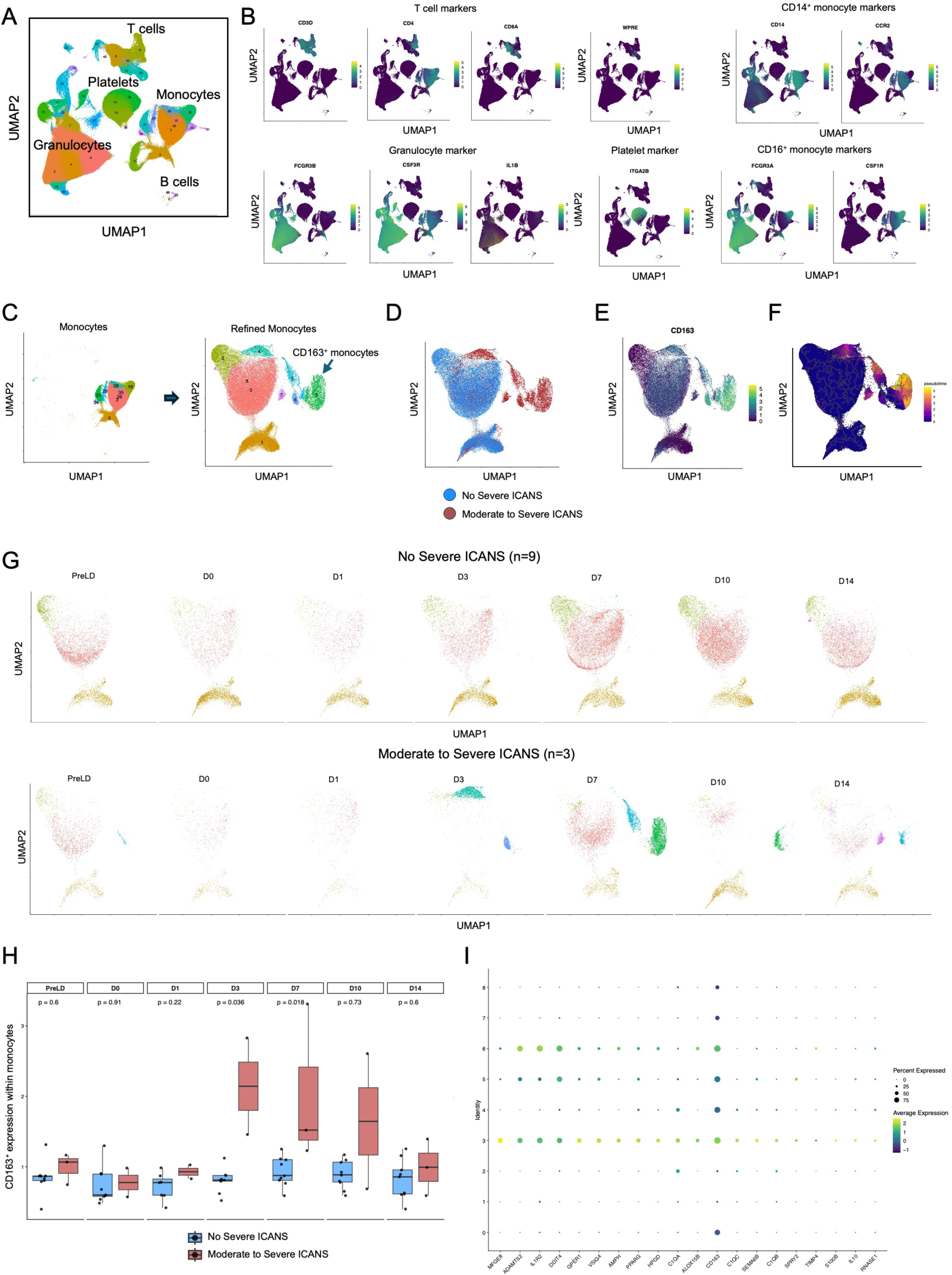
Transcriptomic profiling identifies a CD163⁺ monocyte population enriched for immunoregulatory and wound-healing programs in moderate-to-severe ICANS. A. UMAP visualization of immune cell populations resolved from fixed whole-blood samples using the GEM-X Flex V2 probe set B. Expression of lineage-defining markers used to annotate major immune populations, including T cells, classical monocytes, nonclassical monocytes, granulocytes, and platelets. C. UMAP visualization of reclustered monocyte populations following targeted selection of monocyte cells for enhanced resolution. D. Distribution of monocyte populations stratified by patients with non-severe ICANS and moderate-to-severe ICANS. E. Feature plot showing CD163 expression across monocyte populations. F. Monocyte pseudotime trajectory analysis with root cells defined as pre-lymphodepletion (PreLD) monocytes. G. Temporal distribution of monocyte populations across sample collection timepoints in patients with non-severe and moderate-to-severe ICANS. H. Average CD163 expression across all monocytes stratified by ICANS severity and collection timepoint. I. Top 20 differentially expressed genes defining Monocyte-3. Dot size indicates the proportion of cells expressing each gene, and color intensity represents scaled average gene expression.

Within the monocyte compartment, we identified a population present exclusively in a patient who developed moderate to severe ICANS **(Figure 5D).** This population, termed Monocyte-3, was distinguished by elevated CD163 expression relative to other monocyte subsets **(Figure 5E).** Pseudotime analysis positioned CD163⁺ monocytes at the terminal end of the inferred monocyte trajectory, distal to Pre-LD monocytes, indicating substantial transcriptional remodeling relative to baseline monocyte populations **(Figure 5F).** Notably, Monocyte-3 was first detected at the Day 7 sampling timepoint in a patient who developed grade 3 ICANS, whose peak neurotoxicity occurred on Day 6. **(Figure 5G).** Consistent with these findings, analysis of CD163 expression across all monocytes revealed increased CD163 expression in patients who developed moderate to severe ICANS as early as Day 3 following CAR T-cell infusion **(Figure 5H).** Elevated CD163 expression persisted through Days 7 and 10 before declining toward baseline levels by Day 14.

Differential gene expression analysis revealed that Monocyte-3 was enriched for wound-healing, immunoregulatory, and anti-inflammatory transcriptional programs. In addition to CD163, top differentially expressed genes included *IL1R2, PPARG,* the complement genes *C1QA, C1QB,* and *C1QC, VSIG4, NFKBIA, and IL10* **(Figure 5I, Supplemental Figure 15A).** Genes overexpressed within this population were further enriched for pathways associated with detoxification and cellular responses to toxic substances, including detoxification, cellular detoxification, response to toxic substance, and cellular response to toxic substance (**Supplemental Figure 15B**). Gene set enrichment analysis also identified enrichment of corticosteroid and glucocorticoid response pathways within CD163⁺ monocytes (**Supplemental Figure 15C**). Although expression of the glucocorticoid receptor gene *NR3C1* was not elevated in this population, CD163⁺ monocytes exhibited increased *FKBP5* expression a canonical glucocorticoid-responsive gene (**Supplemental Figure 15D**)^24^.

To further explore functional properties, we applied CellChat across our broad cellular clustering to infer intercellular communication networks (**Supplemental Figure 16A-B)**^25^. Based on ligand–receptor expression patterns, CD163⁺ monocytes were identified as a probable source of IL-10 signaling to multiple cellular populations, supporting a potential immunoregulatory role **(Supplemental Figure 16C-E).**

Together, these orthogonal single-cell transcriptomic analyses independently validated the emergence of CD163⁺ monocytes in patients who develop moderate-to-severe ICANS. These cells exhibited a transcriptionally distinct state characterized by immunoregulatory, wound-healing, and glucocorticoid-responsive programs and were predicted to participate in IL-10–mediated intercellular communication. These findings support the expansion of CD163⁺ monocytes as a reproducible feature of severe ICANS and suggest that they represent a highly remodeled monocyte population that emerges in the setting of neurotoxicity, potentially in response to inflammatory signals, therapeutic interventions, or a combination of both.

### Transcriptomic analysis highlights presence of CD177^hi^ CD10^low^ neutrophil population associated with moderate to severe ICANS\

In parallel, we performed high-resolution reclustering of *FCGR3B-*enriched granulocyte populations **(Figure 6A).** This analysis identified multiple granulocyte subsets, including populations that emerged specifically in patients with moderate to severe ICANS **(Figure 6B).** We observed that granulocytes originating from patients with moderate to severe ICANS were enriched for *IL1R2* and *IL18R1* expression **(Figure 6C).** Consistent with findings in CD163⁺ monocytes, granulocyte populations from patients who developed moderate to severe ICANS also exhibited elevated FKBP5 expression **(Supplemental Figure 17).** One such population, Granulocyte 7, was characterized by low expression of CD10 (*MME*), consistent with the CD10⁻ neutrophil population identified by mass cytometry **(Figure 6C).** Additionally, Granulocyte-7 emerged at the Day 7 collection timepoint, corresponding closely with peak grade 3 ICANS onset on Day 6 following CAR T cell infusion in this patient **(Figure 6D).**

**Figure 6.**
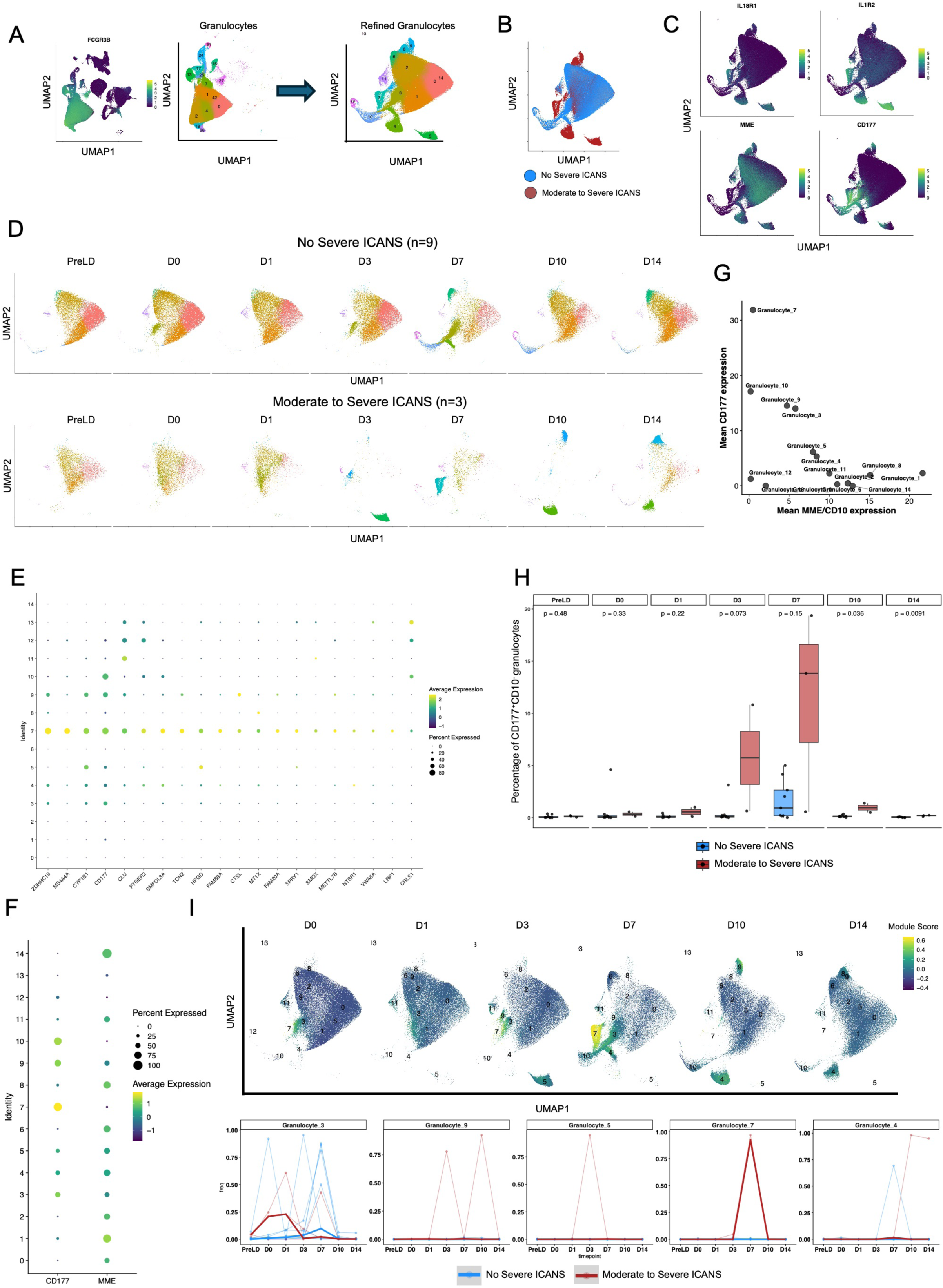
Transcriptomic validation identifies CD177^hi^CD10^low^ neutrophils enriched in patients with moderate-to-severe ICANS. A. Feature plot of FCGR3B expression across all immune populations identified by GEM-X Flex profiling and UMAP visualization of granulocyte populations following targeted selection and reclustering for enhanced resolution. B. Distribution of granulocyte populations stratified by patients with non-severe ICANS and moderate-to-severe ICANS. C. Feature plots showing expression of *IL18R1*, *IL1R2*, *CD177*, and *MME* (CD10) across granulocyte populations. D. Temporal distribution of granulocyte populations across sample collection timepoints in patients with non-severe and moderate-to-severe ICANS. E. Top 20 differentially expressed genes defining Granulocyte-7. Dot size indicates the proportion of cells expressing each gene, and color intensity represents scaled average gene expression. F. Dot plot showing *CD177* and *MME* (CD10) expression across granulocyte populations. Dot size indicates the proportion of cells expressing each gene, and color intensity represents scaled average gene expression. G. Relationship between average *CD177* and average *MME* (CD10) expression across granulocyte populations H. Frequency of CD177^hi^CD10^low^ granulocytes across longitudinal sampling timepoints stratified by ICANS severity. I. Top: Granulocyte-7 module scores across all granulocytes over the course of CAR T cell therapy. Bottom: Longitudinal abundance of granulocyte populations exhibiting Granulocyte-7–like transcriptional signatures in patients with non-severe and moderate-to-severe ICANS. Lines connect serial samples collected from the same patient, and bold lines indicate group medians.

Because *CD101* was not included in the 10X FLEX APEX probe set, we sought alternative markers to define this CD10^low^ granulocyte population. Differential expression analysis identified *CD177* as a distinguishing marker for our *MME* granulocyte population of interest **(Figure 6E**). CD177 surface expression is of particular interest given its role as a neutrophil activation marker and its function as a receptor for PECAM-1^26^, which has a key role in regulating neutrophil extravasation during inflammation.

Based on prior studies demonstrating elevated CD177:CD10 ratios in inflammatory conditions such as sepsis, we evaluated whether this metric could be applied to ICANS^27^. Granulocyte 7 exhibited the highest *CD177* expression and lowest MME expression among granulocyte subsets **(Figure 6F,G).** Across all granulocyte populations, the *CD177:CD10* ratio was elevated in moderate to severe ICANS cases as early as Day 3 post-infusion and remained persistently elevated relative to non-severe cases through the final collection timepoint (Day 14) **(Figure 6H).**

Because Granulocyte-7 was detected in only 3 of 13 patients, we next sought to identify a broader transcriptomic profile that could overcome limitations associated with rare cluster-based analyses. To accomplish this, we generated a gene module derived from the top 20 differentially expressed genes enriched in Granulocyte-7 and projected this signature across all granulocyte clusters at each collection timepoint **(Figure 6I).** This analysis revealed that multiple granulocyte populations shared transcriptional features with Granulocyte-7, particularly during periods of peak abundance in patients who developed severe ICANS. Across longitudinal CAR T cell therapy timepoints, granulocyte populations exhibiting the highest Granulocyte-7 module scores were consistently enriched in severe ICANS cases.

At Day 0, Granulocyte-3 showed the highest Granulocyte-7 module expression among granulocyte populations and was enriched in patients with moderate to severe ICANS. By Day 3, this transcriptional program was most strongly enriched within Granulocyte-9 cells, which were likewise enriched in moderate to severe ICANS cases. At later post-infusion timepoints, including Days 7 and 10, elevated Granulocyte-7-like transcriptional signatures were observed within Granulocyte-7 and Granulocyte-4 populations, respectively, both of which were enriched in moderate to severe ICANS patients.

Together, these findings suggest that transcriptomic programs associated with immature or inflammatory granulocyte states are distributed across multiple neutrophil populations and correlate with ICANS severity throughout the course of CAR T cell therapy.

Collectively, these transcriptomic analyses independently validated the emergence of immature granulocyte populations in patients who develop moderate-to-severe ICANS. These cells were characterized by reduced *CD10* expression, increased *CD177* expression and CD177:CD10 ratios, elevated IL1R2 and *IL18R1* expression, and glucocorticoid-responsive transcriptional programs. Importantly, the Granulocyte-7 transcriptional signature extended beyond a single rare cluster and was observed across multiple granulocyte populations enriched in severe ICANS, supporting the existence of a broader immature inflammatory neutrophil state associated with CAR T-cell–related neurotoxicity.

## DISCUSSION

In this study, we employed a whole-blood–compatible multimodal platform to longitudinally profile peripheral immune cell populations and serum proteins associated with moderate-to-severe ICANS in patients receiving CD19 CAR T-cell therapy. By integrating mass cytometry, serum proteomics, and single-cell transcriptomics, we identified cellular and soluble biomarkers associated with neurotoxicity within a three-day window surrounding peak ICANS onset. To enhance temporal resolution and capture dynamic immune changes, sample collection timepoints were aligned relative to each patient’s peak ICANS or CRS diagnosis. A major strength of our approach was the use of SMART Tube Proteomic Stabilizer, which enabled preservation and high-dimensional profiling of typically fragile granulocyte populations that are often underrepresented in conventional single-cell workflows. This strategy allowed high-resolution characterization of neutrophil heterogeneity throughout CAR T-cell therapy and revealed dynamic changes associated with neurotoxicity.

Using this approach, we found that neutrophils from patients undergoing CD19 CAR T-cell therapy were phenotypically distinct from those of healthy donors even prior to infusion. Although the mechanisms underlying these differences remain unclear, prior chemotherapy exposure and administration of G-CSF before CAR T-cell therapy have been proposed to contribute to the release of atypical neutrophil populations from the bone marrow into circulation^28^. However, we did not observe significant correlations between cumulative filgrastim exposure and the abundance of CD177^hi^CD10^low^ neutrophils in our cohort **(Supplemental Figure 10E-F),** suggesting that G-CSF administration alone is unlikely to fully explain the emergence of these populations.

Importantly, longitudinal profiling revealed expansion of immature CD10^low^ neutrophil subsets in patients who developed moderate-to-severe ICANS. CD10 has previously been characterized as a marker of neutrophil maturity and is reduced in patients with sepsis and other inflammatory conditions.^29^. Consistent with these observations, we found reduced CD10 expression within neutrophils from patients with moderate-to-severe ICANS at peak toxicity. In a smaller validation cohort, we further observed elevated CD177 expression within this same population and identified CD177 as a defining differentially expressed gene. CD177 has been reported among the most highly upregulated neutrophil-associated genes in severe COVID-19 and other inflammatory syndromes, while the *CD177:CD10* gene expression ratio has emerged as a marker of neutrophil activation and disease severity in sepsis.^30,31^. Together, these findings suggest that reduced CD10 expression coupled with elevated CD177 expression identifies a population of immature, activated neutrophils associated with severe ICANS.

It was also noteworthy that CD177⁺ granulocytes exhibited elevated expression of FKBP5. FKBP5 has been implicated in promoting neuroinflammation through neutrophil extracellular trap (NET) formation in ischemic stroke, and excessive NETosis has been described in numerous inflammatory conditions including COVID-19^32,33^. Given the activated phenotype of CD177⁺ granulocytes observed in our cohort, elevated FKBP5 expression raises the possibility that these cells contribute to ICANS-associated inflammation through NET-associated mechanisms. Although further mechanistic studies are required, these observations suggest that dysregulated neutrophil activation may represent an important component of ICANS pathogenesis.

Our observations are consistent with emerging evidence that myeloid cells play an important role in shaping outcomes following CD19 CAR T-cell therapy. While early studies largely focused on CAR T-cell expansion, persistence, and functional state, more recent work has identified monocyte and neutrophil populations associated with treatment response and survival^13,14^. Monocytic transcriptional programs characterized by *MS4A4A*, *CD86*, *CD163*, and *SIGLEC5* expression within leukapheresis products have been associated with inferior progression-free survival, while immature CD10⁻ neutrophils have similarly been linked to adverse clinical outcomes following CAR T-cell therapy^34,35^. Together with our findings, these studies support a broader role for myeloid dysregulation in shaping both therapeutic responses and treatment-related toxicities.

Alongside the expansion of immature neutrophil populations, we observed a significant enrichment of CD163⁺ monocytes at peak ICANS among patients who developed moderate-to-severe neurotoxicity. CD163⁺ monocyte abundance positively correlated with cumulative dexamethasone exposure from CAR T-cell infusion through peak ICANS diagnosis, consistent with prior reports demonstrating glucocorticoid-mediated induction of CD163 expression on monocytes and macrophages **(Supplemental Figure 5A-B)**^36^. Supporting this association, CD163⁺ monocytes identified in our validation cohort exhibited enrichment of glucocorticoid-response pathways. However, whether these cells arise primarily as a consequence of ICANS-associated inflammation, corticosteroid exposure, or a combination of both remains unclear. We did not observe significant associations between CD163⁺ monocyte abundance and cumulative methylprednisolone or filgrastim exposure administered from Day 0 through peak ICANS. However, these analyses may have been underpowered due to the relatively small number of patients who received each treatment (n = 9 and n = 17, respectively). Similarly, CD10^low^ neutrophil abundance was not significantly associated with cumulative dexamethasone or methylprednisolone exposure **(Supplemental Figure 10A–D).**

Among serum proteins quantified at peak ICANS, ST2 and IL-2RA demonstrated the strongest correlations with CD163⁺ monocyte abundance **(Figure 4F).** Notably, soluble ST2, CD163, and IL-2RA are established biomarkers of hemophagocytic lymphohistiocytosis (HLH), another inflammatory toxicity associated with CAR T-cell therapy, and are similarly elevated in other hyperinflammatory conditions including SARS-CoV-2 infection^37,38^. These observations suggest shared features of immune dysregulation across inflammatory syndromes characterized by excessive immune activation^39,40^. Previous studies have shown that elevated soluble CD163 following traumatic injury originates from peripheral monocytes and contributes to immune regulation through crosstalk with lymphocyte populations. Consistent with this literature, our findings suggest that CD163⁺ monocytes may represent an immunoregulatory response that emerges following the onset of severe inflammation^41^.

Additional evidence supporting an immunoregulatory phenotype was observed within the transcriptional profile of these cells. CD163⁺ monocytes exhibited elevated expression of VSIG4, NFKBIA, and PPARG relative to other monocyte populations. VSIG4 has previously been linked to lymphoma-associated HLH and functions as a negative regulator of macrophage activation, while NFKBIA and PPARG are established suppressors of inflammatory signaling pathways.^42,43,44^. Together, these findings suggest that CD163⁺ monocytes may represent a compensatory myeloid population that emerges in response to severe inflammation and contributes to immune regulation during ICANS. Future studies quantifying soluble CD163 and VSIG4 in serum may help determine whether these factors serve as biomarkers of myeloid activation, inflammatory resolution, treatment-induced effects, or toxicity-associated immune regulation following CAR T-cell therapy.

Pseudotime analysis further revealed that the CD163⁺ monocyte population, which peaked at Day 7, occupied the terminal end of the inferred monocyte trajectory and was transcriptionally most distinct from baseline Pre-LD monocytes. In contrast, monocytes collected at Days 10 and 14 occupied states that more closely resembled baseline populations. These findings suggest that monocyte activation reaches its greatest magnitude near peak ICANS before progressively resolving as the immune compartment returns toward homeostasis.

Furthermore, while we observed a positive correlation between cumulative dexamethasone dose from CAR T cell infusion through peak ICANS diagnosis and CD163⁺ monocyte abundance, this association does not establish causality. Thus, it remains unclear whether these CD163⁺ monocytes arise primarily as part of the endogenous inflammatory response to ICANS, as a consequence of corticosteroid exposure administered to mitigate toxicity, or through a combination of both processes.

A notable finding across both neutrophil and monocyte compartments was the elevated expression of IL-1R2.

IL-1R2 functions as a decoy receptor that attenuates IL-1 signaling by competing with IL-1R1 for ligand binding and sequestering IL-1RAP, thereby preventing formation of a signaling-competent receptor complex.^45,46^. IL1R2 expression is induced by glucocorticoids such as dexamethasone and can also be upregulated by IL-33 as part of a negative feedback mechanism that limits inflammatory signaling.^47,48^. Given that IL-33 is primarily released from endothelial and epithelial cells and signals through ST2, these findings suggest the presence of a coordinated regulatory axis involving IL-33, ST2, and IL-1R2^49^. We speculate that activation of this network reflects a compensatory immunomodulatory response to inflammation and may contribute to shaping ICANS severity following CAR T-cell therapy.

In addition to the emergence of CD163⁺ monocytes, we observed a reduced proportion of peripheral non-classical CD16⁺ monocytes in patients who developed moderate-to-severe ICANS, particularly following peak toxicity. This observation is consistent with reports demonstrating inverse correlations between circulating CD16⁺ monocytes and disease severity in COVID-19^50^. During acute inflammatory responses, non-classical monocytes migrate from the peripheral circulation into inflamed tissues, where they contribute to local immune responses^51^. In severe COVID-19, these cells accumulate within distal lung tissues and are thought to contribute to disease pathology.^52^. Similarly, previous studies have demonstrated the ability of CD16⁺ monocytes to traverse the brain microvascular endothelial barrier and enter the central nervous system under inflammatory conditions^53^. Thus, reduced peripheral CD16⁺ monocyte abundance in patients with severe ICANS may reflect trafficking of these cells into the central nervous system or associated vascular compartments. Consistent with this possibility, blood count analyses revealed an overall reduction in circulating monocyte counts among patients with moderate-to-severe ICANS, suggesting altered monocyte trafficking or impaired recovery following lymphodepletion.

Several limitations should be considered when interpreting our findings. First, sample sizes were modest for several analyses, particularly those integrating cellular and serum measurements at peak ICANS, limiting statistical power and the ability to detect smaller effects. Second, the observational nature of this study precludes conclusions regarding causality. Whether the emergence of CD163⁺ monocytes and immature neutrophils directly contribute to ICANS pathogenesis or instead represents a downstream consequence of inflammatory processes or therapeutic interventions remains unresolved. Third, although our findings were supported by an independent validation cohort, larger cohorts will be required to establish the robustness and generalizability of these observations across CAR T-cell products and disease settings.

Future studies incorporating higher-frequency longitudinal sampling, mechanistic model systems, and functional characterization of neutrophil and monocyte subsets will be necessary to define their temporal relationships to ICANS onset and determine whether they play causal roles in disease pathogenesis. Additional mass cytometry studies in independent CAR T-cell cohorts and expanded single-cell transcriptomic analyses may reveal additional myeloid populations and molecular pathways involved in neurotoxicity. Mechanistic investigations using co-culture systems, blood–brain barrier models, and in vivo approaches will be particularly important for defining how peripheral myeloid cells influence central nervous system inflammation following CAR T-cell therapy.

Overall, our study demonstrates the utility of longitudinal whole-blood immune profiling for identifying cellular and soluble correlates of CAR T-cell–associated toxicities. The identification of immature CD10⁻CD177⁺ neutrophils and immunoregulatory CD163⁺ monocytes as features of moderate-to-severe ICANS provides new insight into the myeloid mechanisms associated with neurotoxicity. These findings raise the possibility that therapeutic strategies targeting neutrophil activation, trafficking, maturation, or effector functions could mitigate neuroinflammatory processes underlying ICANS. Furthermore, longitudinal monitoring of these populations may enable earlier identification of patients at increased risk for severe neurotoxicity, creating opportunities for biomarker-guided intervention before the onset of overt neurological symptoms. While further mechanistic studies are required to establish causality, our findings nominate immature neutrophils and CD163⁺ monocytes as candidate biomarkers and potential therapeutic targets for reducing CAR T-cell therapy–associated neurotoxicity.

## METHODS

### Study Design and Participants profiled with Mass Cytometry

Thirty-three patients with relapsed and/or refractory B cell malignancies underwent CD19 CAR T cell therapy. The patients received commercial Yescarta (n=18), Tecartus (n=3), Kymriah (n=6), or Breyanzi (n=6) infusions. Of the 33 patients, 18 achieved complete remission by day 28 post CAR T cell infusion, while 15 experienced only partial response or disease progression. To define and characterize cellular populations and soluble mediators associated with moderate to severe ICANS in CD19 CAR T cell therapy we established a whole blood and serum collection protocol for the 33 patients. Whole blood samples and serum were collected at multiple timepoints: prior to CAR T cell infusion, at the onset of adverse event, at the time of peak toxicity, and at toxicity resolution. Blood samples were fixed with SMARTube Proteomic Stabilizer buffer and cryopreserved to maintain neutrophil and eosinophil structural integrity for downstream CyTOF analysis. We designed a 40-marker antibody mass cytometry panel incorporating lineage, activation, memory, proliferation, phosphorylation, and migration markers to comprehensively characterize the diverse immune populations in CD19 CAR T cell therapy.

### Mass Cytometry whole blood fixation and cellular staining

Whole blood was mixed with SMARTube Proteomic Stabilizer buffer in 1:1.4 ratio and left at room temperature for 10 minutes prior to storage in -80 in 2ml aliquots. On the day of cellular staining, the whole blood and fixation aliquots were thawed on ice for 15 minutes before addition of 2 ml SMARTtube Thaw-Lyse buffer and incubated for an additional 10 minutes to deplete red blood cells. 3 rounds of SMARTtube Thaw-Lyse washes were performed afterwards prior to transferring cells to a 96 well plate and incubation in a 9:1 Fc Receptor Blocking solution and 200 U Heparin mixture at 4 C for 20 minutes. Cells were washed and resuspended in cytometry buffer (PBS + 0.02% sodium azide + 5% fetal calf serum). Following a wash with cytometry buffer, cells were incubated with PE-conjugated antibody against TCRgd on ice for 15 minutes. Afterward, cells were washed with cytometry buffer and stained with a cocktail of metal-conjugated surface marker antibodies at titrated concentrations and incubated on ice for 30 minutes. Following surface marker staining, cells were washed in cytometry buffer twice prior to permeabilization with methanol on ice for 10 minutes. Following permeabilization and two washes with cytometry buffer, cells were incubated with a cocktail of metal-conjugated antibodies against intracellular transcription factors at room temperature for 30 minutes. An additional two rounds of wash with cytometry buffer followed before incubating with 2% paraformaldehyde and DNA intercalator (1:4000) and storing at 4C until the day of CyTOF acquisition.

### CyTOF acquisition

Over the span of several months, all samples were run on the CyTOF in 18 batches, typically composed of the repeated measures from 1-2 patients, as well as a repeated healthy donor control.

Each batch, cells were washed 3× with cytometry buffer and resuspended in 10% EQ beads (Catalog #201078) in MaxPar Cell Acquisition Solution prior to CyTOF processing.

Following CyTOF processing, the signal of each parameter was normalized based on EQ beads with Matlab software (https://github.com/nolanlab/bead-normalization/releases/tag/v0.3. Based on CD66b, cellular populations were gated and exported for clustering analysis using customized R scripts.

### Clustering analysis

Repeated clustering at the single cell level – typically necessary to identify and factor out technical noise, or putative batch effect -- is computationally intractable at the scale of this study. Hence, to arrive at globally defined clusters across all batches, we first ‘over’-clustered within each batch by clustering then subclustering each cluster^54^. Next we computed the centroids of all clusters in each batch, then (meta)clustered these centroids (or ‘‘metacells’’ or “supercells”). At each stage we used the Rphenograph algorithm, with Leiden clustering of shared nearest-neighbor graphs^55^.

To address moderate drift across batches in some parameters, we aligned the repeated control median values across batches for each parameter and applied the corresponding batch- and parameter-specific shifts to the samples before computing metacells for global metaclustering^56^. We used Uniform Manifold Approximate Projection (UMAP) for dimension reduction and data visualization^57^.

### Generalized Estimating Equation

To determine whether clinical grouping by CRS and/or ICANS severity held marginal associations over post-infusion timepoints, for any specific clusters or serum proteins, we applied Generalized Estimating Equations using the R package geepack^58,59^. We used a first-order autoregressive correlation structure (corstr == “ar1”), with non-uniform patient sample timepoints centered according to each patient’s peak ICANS date (set to 0). Multiple hypothesis correction was employed with a Benjamini-Hochberg FDR < .05 for statistical significance.

### Meso-Scale Discovery Assay

To minimize freeze–thaw cycles, serum samples were thawed once, aliquoted into 96-well plates at the required volume for each assay, and then re-frozen until the day of the assay. Given the large number of samples, each assay was run in two batches. Pooled healthy donor serum spiked with the top standard (STD1) was included on all plates as the inter-plate control, since many pro-inflammatory analytes are undetectable in healthy serum. All assays used in the analysis had an inter-plate control coefficient of variation (CV) between 3–16%.

Samples were run in technical duplicate, and the average concentration (pg/ml) for each analyte was interpolated using a weighted four-parameter logistic (4-PL) fit curve defined by 10 non-zero standards using the Meso Scale Discovery software. Assays were performed on either the Meso Scale Discovery (MSD) platform or by traditional ELISA, according to the manufacturer’s instructions, with sample dilutions listed in Table 1.

The lower limit of detection (LLOD) was calculated as the average blank (or raw signal in the sample dilution matrix) plus 2.5 standard deviations. Values above the blank but below the LLOD were interpolated by MSD software and left unchanged. Similarly, values above the curve fit of the top standard (upper limit of detection, ULOD) were interpolated by MSD software and left unchanged. Most cytokines fell within detectable ranges, if an analyte had < 20% of samples in detectable ranges those analytes were removed from further analysis **(Supplemental Table 3).** For transparency, both the LLOD and ULOD are indicated on graphs where values fall below or above these ranges.

Missing cytokine values due to signals below the curve fit or below the average blank were replaced with the re-scaled LLOD (based on sample dilution) to avoid zeros in the dataset and enable log transformation.

### Immune Module Analysis

Effect sizes for each feature of interest were calculated. Subsequently, dimensionality reduction was performed via PCA to group our features of interest into immune modules^23^. We used random forest to categorically classify each immune module as a predictor of moderate to ICANS and rank immune modules based on variable importance, selecting the model with the lowest out-of-bag error as a representative model.

### Validation Cohort Study Design and Participants

Twelve patients with relapsed and/or refractory B-cell malignancies underwent CD19 CAR T-cell therapy with the commercial product Breyanzi. To enable longitudinal detection of CAR T cells, infusion products were manufactured using a lentiviral vector containing a Woodchuck Hepatitis Virus Posttranscriptional Regulatory Element (WPRE) sequence flanking the CAR transgene^60^.Of these, three patients developed moderate to severe ICANS. Whole blood was collected longitudinally at pre-lymphodepletion (PreLD) and on Days 0, 1, 3, 7, 10, and 14 following CAR T cell infusion. Blood samples were fixed in freshly prepared 4% formaldehyde diluted in PBS and stored at 4°C for one week prior to processing. Fixed samples were subsequently mixed with 50% glycerol and stored at −80°C until red blood cell depletion. Red blood cells were depleted using EasySep RBC Depletion Reagent (STEMCELL Technologies). Following red blood cell depletion, samples were resuspended in 10X Enhancer supplemented with 50% glycerol and stored at −80°C. Fixed leukocytes were then hybridized with transcript-specific probes using GEM-X FLEX v2 Human Transcriptome Probe Kit in a 96-well plate prior to the addition of sample-specific multiplexing barcodes. All 96 samples were combined to minimize multiplets and 4 million cells were subsequently processed in four Gel Bead in Emulstions (GEMs) using the 10x Genomics GEM-X Flex v2 kit, according to the manufacturer’s instructions (CG000834 RevA). Resulting libraries were sequenced on a NovaSeqX (Illumina) and subjected to downstream single-cell transcriptomic analysis.

### Statistical Analysis

We designated time bins according to expected timeframes of biological responses, relative to each patient’s peak ICANS diagnosis time, for those with ICANS, or 3 days after infusion for patients without ICANS. When patients have with multiple blood and serum samples collected in a single time bin, the maximum cytokine concentration and the median cluster abundance were used. For each time bin, rank-sum Wilcoxon tests were applied to identify statistically significant differences in median cluster abundances between moderate to severe and non-severe ICANS patient groups. Benjamini-Hochberg multiple test correction was done using FDR < .05 for statistical significance. Firth logistic regression was used to determine statistically significant cytokines associated with moderate to severe ICANS within the peak ICANS window. For the cytokine analyses, an FDR threshold of <0.1 was applied to account for the smaller sample size (n=16) relative to the mass cytometry dataset at peak ICANS diagnosis (n=31). For correlation analysis between cytokine concentrations and cluster abundances, Spearman rank correlation was used.

## AUTHORS DISCLOSURES

J.G. reports personal fees from Bristol Myers Squibb, Kite Pharma, and Cartesian Therapeutics outside the submitted work. J.G. also reports research funding from Sobi, Juno Therapeutics, Celgene, Angiocrine Bioscience, Faron Pharmaceuticals, CARGO Therapeutics, CytoAgents, and Miltenyi Biotec, and service on Independent Data Review Committees for Century Therapeutics and the University of Pennsylvania. J.J.H receives research funding to the institution from Bristol Myers Squibb, BeOne Medicines, and Abbvie and has performed consultancy with AbbVie. S.R.R. is a founder of, advisor to, and inventor on patents licensed to Juno Therapeutics, a Bristol Myers Squibb company. S.R.R. is also a founder of and equity holder in Lyell Immunopharma and has served on advisory boards for Lyell Immunopharma, AstraZeneca, Adaptive Biotechnologies, Outpace Bio, ErVaccine, and Nohla. E.W.N. is a co-founder, advisor and shareholder for ImmunoScape Pte. Ltd. The other authors declare no completing interests. No other disclosures were reported.

## AUTHORS’ CONTRIUBTIONS

**T. Chour:** Conceptualization, Methodology, Validation, Formal Analysis, Investigation, Data Curation, Visualization, Writing – Original Draft, Writing – Review & Editing. **N. Poole:** Investigation, Data Curation, Writing – Review & Editing. **H. MacMillian:** Methodology, Data Curation, Visualization, Writing – Review & Editing. **K. Burleigh:** Investigation, Data Curation. **D.R. Glass:** Methodology. **E.C. Liang:** Data Curation, Visualization **R. Basom:** Data Curation. **B. Webb-Robertson:** Formal Analysis. **K. Stratton:** Formal Analysis. **D. Gratz:** Data Curation. **A.N. Long:** Investigation. **A.E. Elz**: Data Curation. **J. Huang:** Data Curation. **A. Hirayama:** Clinical Investigation. **S.R. Riddell:** Funding Acquisition, Writing – Review & Editing. **J. Gauthier:** Clinical Investigation, Data Curation. **H.H. Gustafson:** Conceptualization, Methodology, Funding Acquisition, Writing – Review & Editing. **E.W. Newell:** Supervision, Conceptualization, Funding Acquisition, Writing – Review & Editing. **S. Simon:** Supervision, Conceptualization, Methodology, Investigation, Data Curation, Funding Acquisition, Writing – Review & Editing.

## ACKNOWLEDGMENTS

T.C. was supported by the Molecular Medicine Training Program through NIH award T32GM095421. N.P. was supported by the NCOR-KUH TL1 Training Program. S.S. was a Special Fellow of the Leukemia & Lymphoma Society Career Development Program (grant 3405-21), supported by The Mark Foundation for Cancer Research. This work was supported by the Fred Hutch Immunotherapy Integrated Research Center (I-IRC), Higgins Research Fund and NIH grants R37CA266777, P01CA018029. The authors acknowledge support for the 8682 Biobank from Swim Across America.

## REFERENCES

1. Schuster, S. J. et al. Chimeric Antigen Receptor T Cells in Refractory B-Cell Lymphomas. N. Engl. J. Med. 377, 2545–2554 (2017).

2. Turtle, C. J. et al. CD19 CAR-T cells of defined CD4+:CD8+ composition in adult B cell ALL patients. J. Clin. Invest. 126, 2123–2138 (2016).

3. Hay, K. A. et al. Kinetics and biomarkers of severe cytokine release syndrome after CD19 chimeric antigen receptor-modified T-cell therapy. Blood 130, 2295–2306 (2017).

4. Gardner, R. A. et al. Preemptive mitigation of CD19 CAR T-cell cytokine release syndrome without attenuation of antileukemic efficacy. Blood 134, 2149–2158 (2019).

5. Gazeau, N. et al. Anakinra for Refractory Cytokine Release Syndrome or Immune Effector Cell-Associated Neurotoxicity Syndrome after Chimeric Antigen Receptor T Cell Therapy. Transplant. Cell. Ther. 29, 430–437 (2023).

6. Frey, N. & Porter, D. Cytokine Release Syndrome with Chimeric Antigen Receptor T Cell Therapy. Biol. Blood Marrow Transplant. J. Am. Soc. Blood Marrow Transplant. 25, e123–e127 (2019).

7. Norelli, M. et al. Monocyte-derived IL-1 and IL-6 are differentially required for cytokine-release syndrome and neurotoxicity due to CAR T cells. Nat. Med. 24, 739–748 (2018).

8. Delgoffe, G. M. et al. The role of exhaustion in CAR T cell therapy. Cancer Cell 39, 885–888 (2021).

9. Fraietta, J. A. et al. Determinants of response and resistance to CD19 chimeric antigen receptor (CAR) T cell therapy of chronic lymphocytic leukemia. Nat. Med. 24, 563–571 (2018).

10. DeFranco, G., Siegler, E. L. & Kenderian, S. S. The Conflicting Role of Myeloid Cells in CAR T-Cell Therapy. Cancer Res. 86, 2581–2591 (2026).

11. McKenna, E. et al. Neutrophils in COVID-19: Not Innocent Bystanders. Front. Immunol. 13, 864387 (2022).

12. Zhang, J. et al. Dysregulation of neutrophil in sepsis: recent insights and advances. Cell Commun. Signal. CCS 23, 87 (2025).

13. Gu, T., Hu, K., Si, X., Hu, Y. & Huang, H. Mechanisms of immune effector cell-associated neurotoxicity syndrome after CAR-T treatment. WIREs Mech. Dis. 14, e1576 (2022).

14. Yang, S. et al. Neutrophil activation and clonal CAR-T re-expansion underpinning cytokine release syndrome during ciltacabtagene autoleucel therapy in multiple myeloma. Nat. Commun. 15, 360 (2024).

15. Qin, J., Wei, F. & Ren, X. Neutrophils in the era of single-cell RNA sequencing: functions and targeted therapies in cancer. Cancer Biol. Med. 20, 903–914 (2024).

16. Zwicklbauer, K. et al. Adapting the SMART tube technology for flow cytometry in feline full blood samples. Front. Vet. Sci. 11, 1377414 (2024).

17. Lee, D. W. et al. ASTCT Consensus Grading for Cytokine Release Syndrome and Neurologic Toxicity Associated with Immune Effector Cells. Biol. Blood Marrow Transplant. J. Am. Soc. Blood Marrow Transplant. 25, 625–638 (2019).

18. Burleigh, K. et al. Low Peripheral Blood Counts and Elevated Proinflammatory Cytokines Signal a Poor CD19 Chimeric Antigen Receptor T-cell Response in Acute Lymphoblastic Leukemia. Transplant. Cell. Ther. 31, 551–564 (2025).

19. Flora, C. et al. Longitudinal plasma proteomics in CAR T-cell therapy patients implicates neutrophils and NETosis in the genesis of CRS. Blood Adv. 8, 1422–1426 (2024).

20. Schulte-Schrepping, J. et al. Severe COVID-19 Is Marked by a Dysregulated Myeloid Cell Compartment. Cell 182, 1419–1440.e23 (2020).

21. Cevering, C. K., Abdel-Azim, H., Khazal, S. J. & Casassa, C. Immune Effector Cell-Associated Neurotoxicity Syndrome: A Practical Overview for the General Neurologist. Neurol. Clin. Pract. 16, e200575 (2026).

22. Moreno-Castaño, A. B. et al. Characterization of the endotheliopathy, innate-immune activation and hemostatic imbalance underlying CAR-T cell toxicities: laboratory tools for an early and differential diagnosis. J. Immunother. Cancer 11, e006365 (2023).

23. Glass, D. R. et al. Multi-omic profiling reveals the endogenous and neoplastic responses to immunotherapies in cutaneous T cell lymphoma. Cell Rep. Med. 5, 101527 (2024).

24. Sato, A., Omichi, R., Maeda, Y. & Ando, M. A glucocorticoid-regulating molecule, Fkbp5, may interact with mitogen-activated protein kinase signaling in the organ of Corti of mice cochleae. Sci. Rep. 15, 7506 (2025).

25. Jin, S., Plikus, M. V. & Nie, Q. CellChat for systematic analysis of cell-cell communication from single-cell transcriptomics. Nat. Protoc. 20, 180–219 (2025).

26. Sachs, U. J. H. et al. The neutrophil-specific antigen CD177 is a counter-receptor for platelet endothelial cell adhesion molecule-1 (CD31). J. Biol. Chem. 282, 23603–23612 (2007).

27. Huang, J. et al. Dynamic CD177/CD10 ratio for infection diagnosis and mortality risk stratification in critically ill patients: a prospective cohort study. EBioMedicine 123, 106100 (2026).

28. Huang, X. et al. Neutrophils in Cancer immunotherapy: friends or foes? Mol. Cancer 23, 107 (2024).

29. Liu, M. et al. Immunoregulatory functions of mature CD10+ and immature CD10- neutrophils in sepsis patients. Front. Med. 9, 1100756 (2022).

30. Lévy, Y. et al. CD177, a specific marker of neutrophil activation, is associated with coronavirus disease 2019 severity and death. iScience 24, 102711 (2021).

31. Demaret, J. et al. Identification of CD177 as the most dysregulated parameter in a microarray study of purified neutrophils from septic shock patients. Immunol. Lett. 178, 122–130 (2016).

32. Li, Z. et al. FKBP5-CCL5 interaction promotes neuroinflammation and neuronal apoptosis in ischemic stroke by regulating the MAPK pathway and enhancing NET formation. Front. Immunol. 16, 1609989 (2025).

33. Zuo, Y. et al. Neutrophil extracellular traps in COVID-19. JCI Insight 5, e138999, 138999 (2020).

34. Carniti, C. et al. Monocytes in leukapheresis products affect the outcome of CD19-targeted CAR T-cell therapy in patients with lymphoma. Blood Adv. 8, 1968–1980 (2024).

35. Zhu, J. et al. Elevated CD10- neutrophils correlate with non-response and poor prognosis of CD19 CAR T-cell therapy for B-cell acute lymphoblastic leukemia. BMC Med. 23, 138 (2025).

36. Svendsen, P., Etzerodt, A., Deleuran, B. W. & Moestrup, S. K. Mouse CD163 deficiency strongly enhances experimental collagen-induced arthritis. Sci. Rep. 10, 12447 (2020).

37. Lovisari, F., Terzi, V., Lippi, M. G., Brioschi, P. R. & Fumagalli, R. Hemophagocytic lymphohistiocytosis complicated by multiorgan failure: A case report. Medicine (Baltimore) 96, e9198 (2017).

38. Rabaan, A. A. et al. SARS-CoV-2 infection and multi-organ system damage: A review. Biomol. Biomed. 23, 37–52 (2023).

39. Gao, Z., Wang, Y., Wang, J., Zhang, J. & Wang, Z. Soluble ST2 and CD163 as Potential Biomarkers to Differentiate Primary Hemophagocytic Lymphohistiocytosis from Macrophage Activation Syndrome. Mediterr. J. Hematol. Infect. Dis. 11, e2019008 (2019).

40. Mostafa, G. A. et al. Up-regulated serum levels of soluble CD25 and soluble CD163 in pediatric patients with SARS-CoV-2. Eur. J. Pediatr. 181, 2299–2309 (2022).

41. O’Connell, G. C. et al. Monocyte-lymphocyte cross-communication via soluble CD163 directly links innate immune system activation and adaptive immune system suppression following ischemic stroke. Sci. Rep. 7, 12940 (2017).

42. Yuan, S. et al. Serum soluble VSIG4 as a surrogate marker for the diagnosis of lymphoma-associated hemophagocytic lymphohistiocytosis. Br. J. Haematol. 189, 72–83 (2020).

43. Li, J. et al. VSIG4 inhibits proinflammatory macrophage activation by reprogramming mitochondrial pyruvate metabolism. Nat. Commun. 8, 1322 (2017).

44. Bouhlel, M. A. et al. PPARgamma activation primes human monocytes into alternative M2 macrophages with anti-inflammatory properties. Cell Metab. 6, 137–143 (2007).

45. Schlüter, T., Schelmbauer, C., Karram, K. & Mufazalov, I. A. Regulation of IL-1 signaling by the decoy receptor IL-1R2. J. Mol. Med. 96, 983–992 (2018).

46. Peters, V. A., Joesting, J. J. & Freund, G. G. IL-1 receptor 2 (IL-1R2) and its role in immune regulation. Brain. Behav. Immun. 32, 1–8 (2013).

47. Re, F. et al. The type II ‘receptor’ as a decoy target for interleukin 1 in polymorphonuclear leukocytes: characterization of induction by dexamethasone and ligand binding properties of the released decoy receptor. J. Exp. Med. 179, 739–743 (1994).

48. Supino, D. et al. Negative Regulation of the IL-1 System by IL-1R2 and IL-1R8: Relevance in Pathophysiology and Disease. Front. Immunol. 13, 804641 (2022).

49. Sheng, F. et al. IL-33/ST2 axis in diverse diseases: regulatory mechanisms and therapeutic potential. Front. Immunol. 16, 1533335 (2025).

50. Gatti, A., Radrizzani, D., Viganò, P., Mazzone, A. & Brando, B. Decrease of Non-Classical and Intermediate Monocyte Subsets in Severe Acute SARS-CoV-2 Infection. Cytom. Part J. Int. Soc. Anal. Cytol. 97, 887–890 (2020).

51. Fahlberg, M. D. et al. Cellular events of acute, resolving or progressive COVID-19 in SARS-CoV-2 infected non-human primates. Nat. Commun. 11, 6078 (2020).

52. Sánchez-Cerrillo, I. et al. COVID-19 severity associates with pulmonary redistribution of CD1c+ DCs and inflammatory transitional and nonclassical monocytes. J. Clin. Invest. 130, 6290–6300 (2020).

53. Ancuta, P., Moses, A. & Gabuzda, D. Transendothelial migration of CD16+ monocytes in response to fractalkine under constitutive and inflammatory conditions. Immunobiology 209, 11–20 (2004).

54. Liu, X. et al. A comparison framework and guideline of clustering methods for mass cytometry data. Genome Biol. 20, 297 (2019).

55. Levine, J. H. et al. Data-Driven Phenotypic Dissection of AML Reveals Progenitor-like Cells that Correlate with Prognosis. Cell 162, 184–197 (2015).

56. Putri, G. H., Howitt, G., Marsh-Wakefield, F., Ashhurst, T. M. & Phipson, B. SuperCellCyto: enabling efficient analysis of large scale cytometry datasets. Genome Biol. 25, 89 (2024).

57. Becht, E. et al. Dimensionality reduction for visualizing single-cell data using UMAP. Nat. Biotechnol. 10.1038/nbt.4314 (2018) doi:10.1038/nbt.4314.

58. Halekoh, U., Højsgaard, S. & Yan, J. The *R* Package **geepack** for Generalized Estimating Equations. J. Stat. Softw. 15, (2006).

59. Liang, K.-Y. & Zeger, S. L. Longitudinal data analysis using generalized linear models. Biometrika 73, 13–22 (1986).

60. Pullarkat, S. et al. qPCR assay for detection of Woodchuck Hepatitis Virus Post-Transcriptional Regulatory Elements from CAR-T and TCR-T cells in fresh and formalin-fixed tissue. PloS One 19, e0303057 (2024).

