## Supplemental Figures for "Coordinated expansion of CD163⁺ monocytes and immature CD177⁺ neutrophils marks severe neurotoxicity after CD19 CAR T cell therapy"

### Supplemental Figure 1

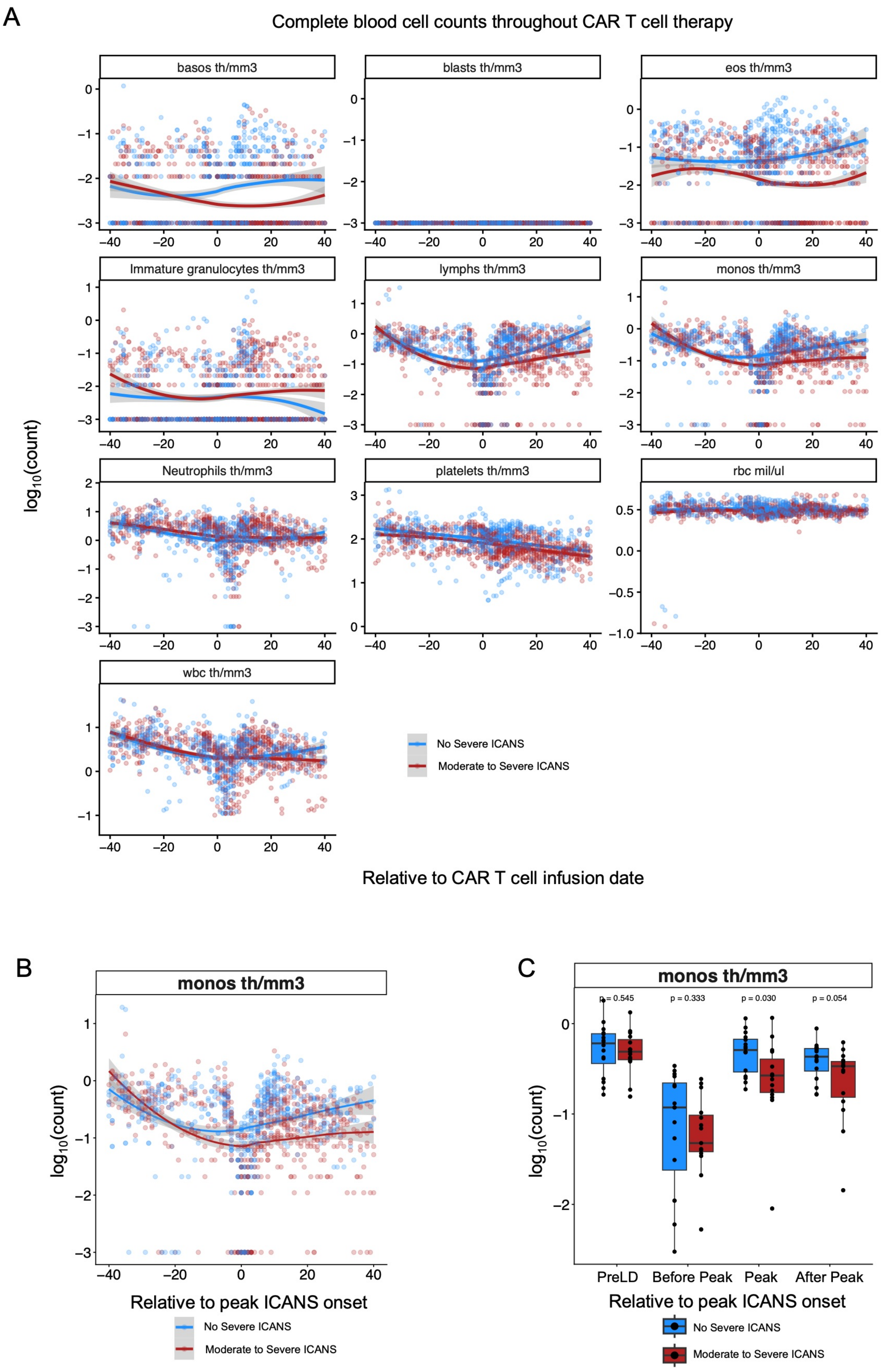

**Supplemental Figure 1. Longitudinal complete blood count dynamics throughout CD19 CAR T cell therapy**

A. Longitudinal trajectories of complete blood count (CBC) parameters measured throughout CD19 CAR T cell therapy and stratified by ICANS severity. Individual points represent patient-matched clinical laboratory measurements collected relative to CAR T cell infusion date. Blue indicates patients who did not develop moderate to severe ICANS, while red indicates patients who developed moderate to severe ICANS. Solid lines represent locally estimated smoothing (LOESS) trends with confidence intervals. CBC parameters shown include basophils, blasts, eosinophils, immature granulocytes, lymphocytes, monocytes, neutrophils, platelets, red blood cells (RBC), and white blood cells (WBC).
B. Longitudinal monocyte counts relative to peak ICANS onset. Timepoint zero represents patient-specific peak ICANS onset.
C. Monocyte counts stratified by ICANS severity across longitudinal ICANS-associated time bins, including pre-lymphodepletion (PreLD), Before Peak ICANS, Peak ICANS, and After Peak ICANS periods. Statistical comparisons were performed using Wilcoxon rank-sum testing.

### Supplemental Figure 2

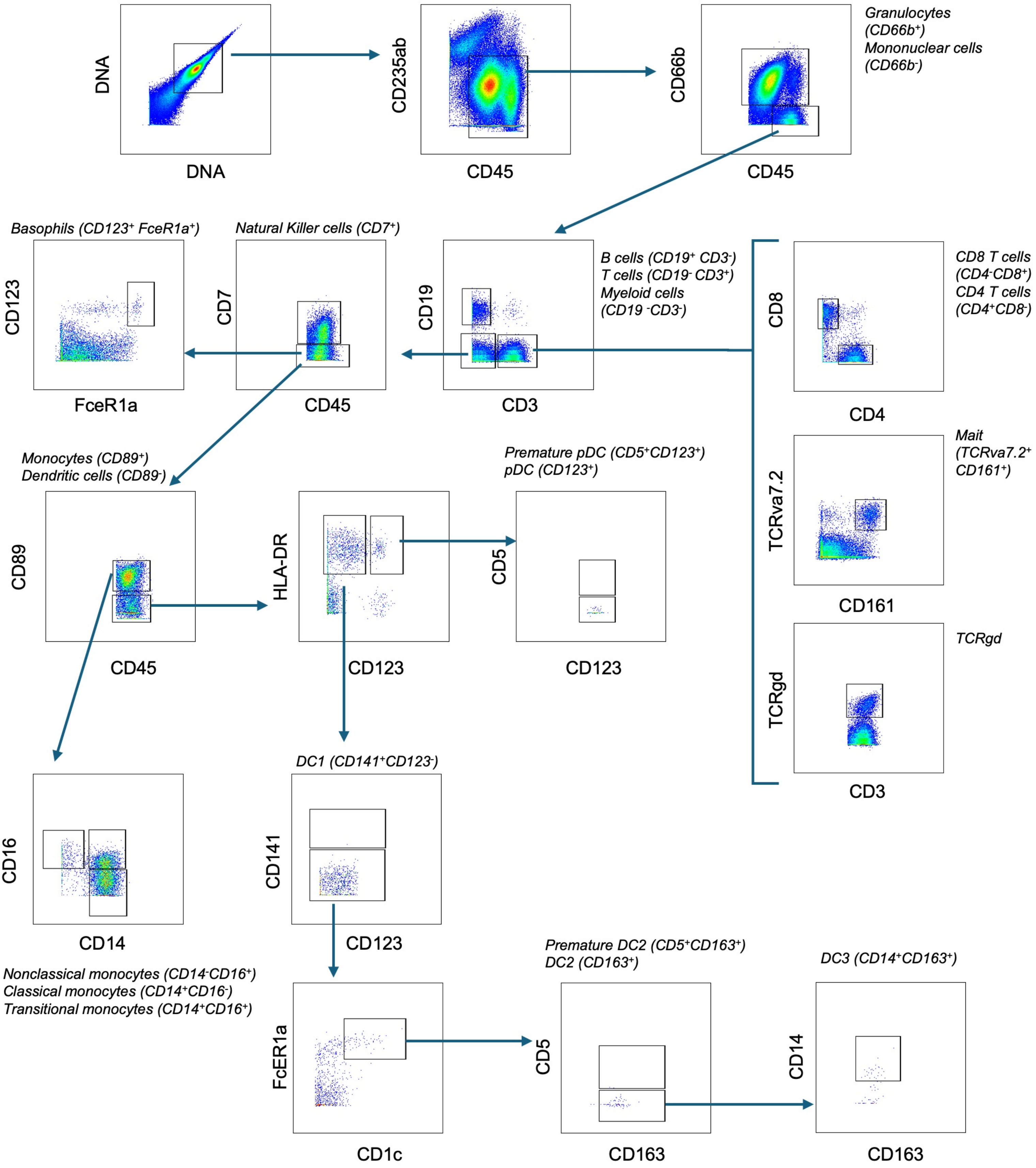

**Supplemental Figure 2. Mass cytometry gating strategy for identification of mononuclear and granulocyte populations.**

Representative biaxial plots illustrating the sequential manual gating strategy used to identify immune populations from fixed whole blood samples analyzed by mass cytometry. Initial gates excluded debris and selected $DNA^{+}CD45^{+}$ leukocytes, followed by separation of $CD66b^{+}$ granulocytes and $CD66b^{-}$ mononuclear cells. Mononuclear cells were further subdivided into basophils, B cells, T cells, TCRγδ T cells, natural killer (NK) cells, mature killer cells, monocytes, and dendritic cell subsets based on canonical lineage marker expression. Monocytes were classified as classical ($CD14^{+}CD16^{-}$), intermediate/transitional ($CD14^{+}CD16^{+}$), and nonclassical ($CD14^{-}CD16^{+}$) subsets. Dendritic cell subsets included DC1 ($CD141^{+}CD123^{-}$), premature plasmacytoid dendritic cells (pDCs; $CD123^{+}CD5^{+}$), premature DC2 ($CD163^{+}$), and DC3 ($CD14^{+}CD163^{+}$) populations. Labels indicate defining marker combinations used for each gated subset.

### Supplemental Figure 3

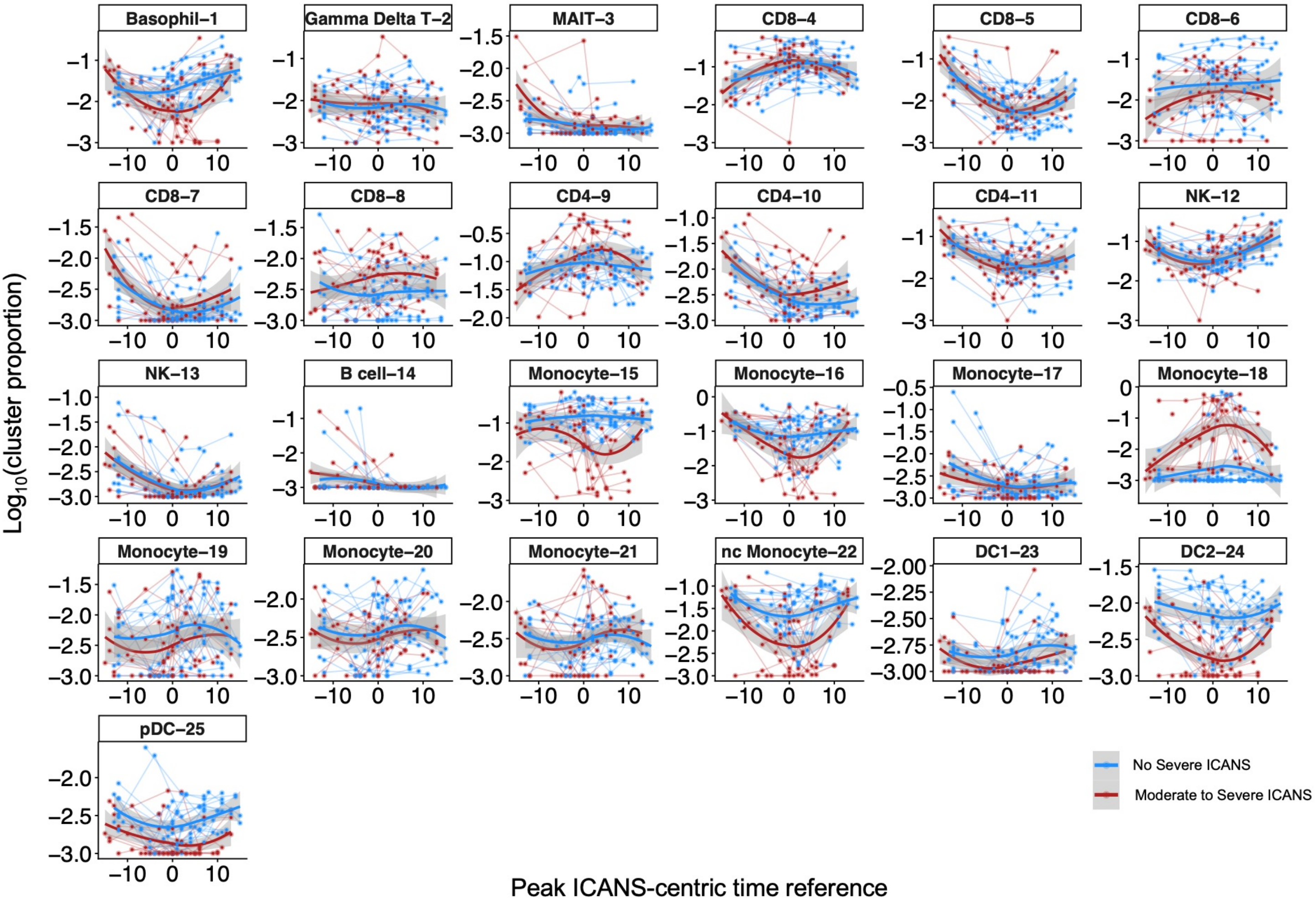

**Supplemental Figure 3. Longitudinal dynamics of mononuclear immune populations relative to peak ICANS onset.**

Longitudinal trajectories of mononuclear population abundances identified by mass cytometry and plotted relative to patient-specific peak ICANS onset. Individual points represent the proportion of each population within the $CD66b^{-}$ mononuclear compartment. Patients with non-severe ICANS are shown in blue and patients with moderate-to-severe ICANS are shown in red. Solid lines represent locally estimated scatterplot smoothing (LOESS) trends with 95% confidence intervals. Populations include T-cell, natural killer (NK) cell, B-cell, monocyte, dendritic cell, basophil, mucosal-associated invariant T (MAIT) cell, and plasmacytoid dendritic cell (pDC) subsets. Population labels correspond to unsupervised clusters identified by high-dimensional mass cytometry analysis.

### Supplemental Figure 4

A

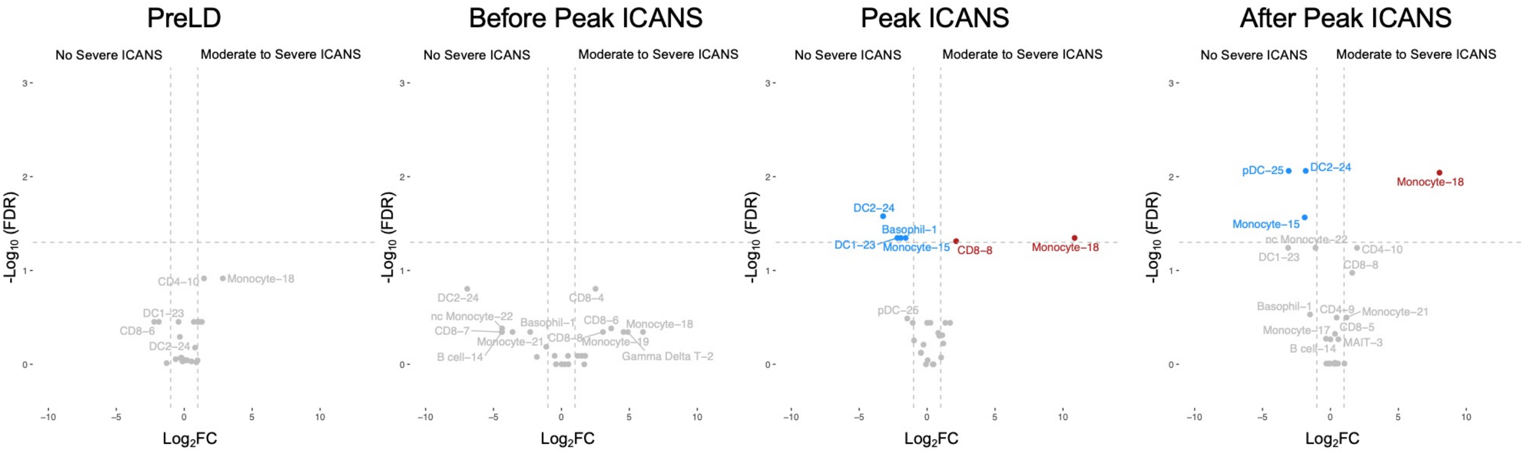


B

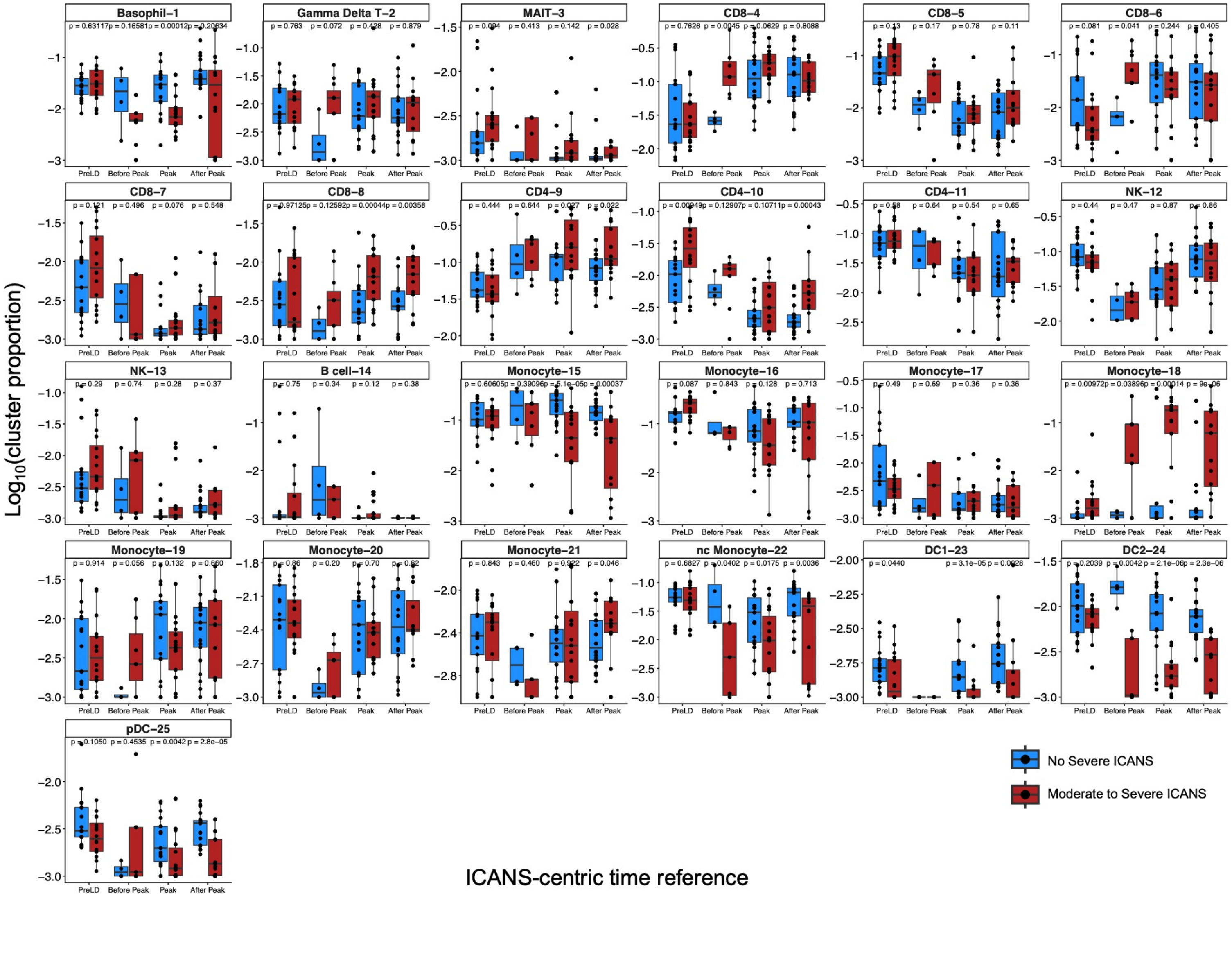

**Supplemental Figure 4. Differential abundance of mononuclear populations associated with ICANS severity.**

A. Volcano plots showing log2 fold changes in mononuclear population abundance between patients with moderate-to-severe ICANS and patients with non-severe ICANS at peak ICANS. Positive values indicate enrichment in moderate-to-severe ICANS, whereas negative values indicate enrichment in non-severe ICANS. Selected populations are annotated, and gray points represent populations that did not reach statistical significance.
B. Relative abundance of mononuclear populations across ICANS-centric time windows, including pre-lymphodepletion (PreLD), Before Peak ICANS, Peak ICANS, and After Peak ICANS, stratified by ICANS severity. Patients with non-severe ICANS are shown in blue and patients with moderate-to-severe ICANS are shown in red. Each point represents an individual patient sample. Statistical comparisons were performed using two-sided Wilcoxon rank-sum tests, with P values indicated.

### Supplemental Figure 5

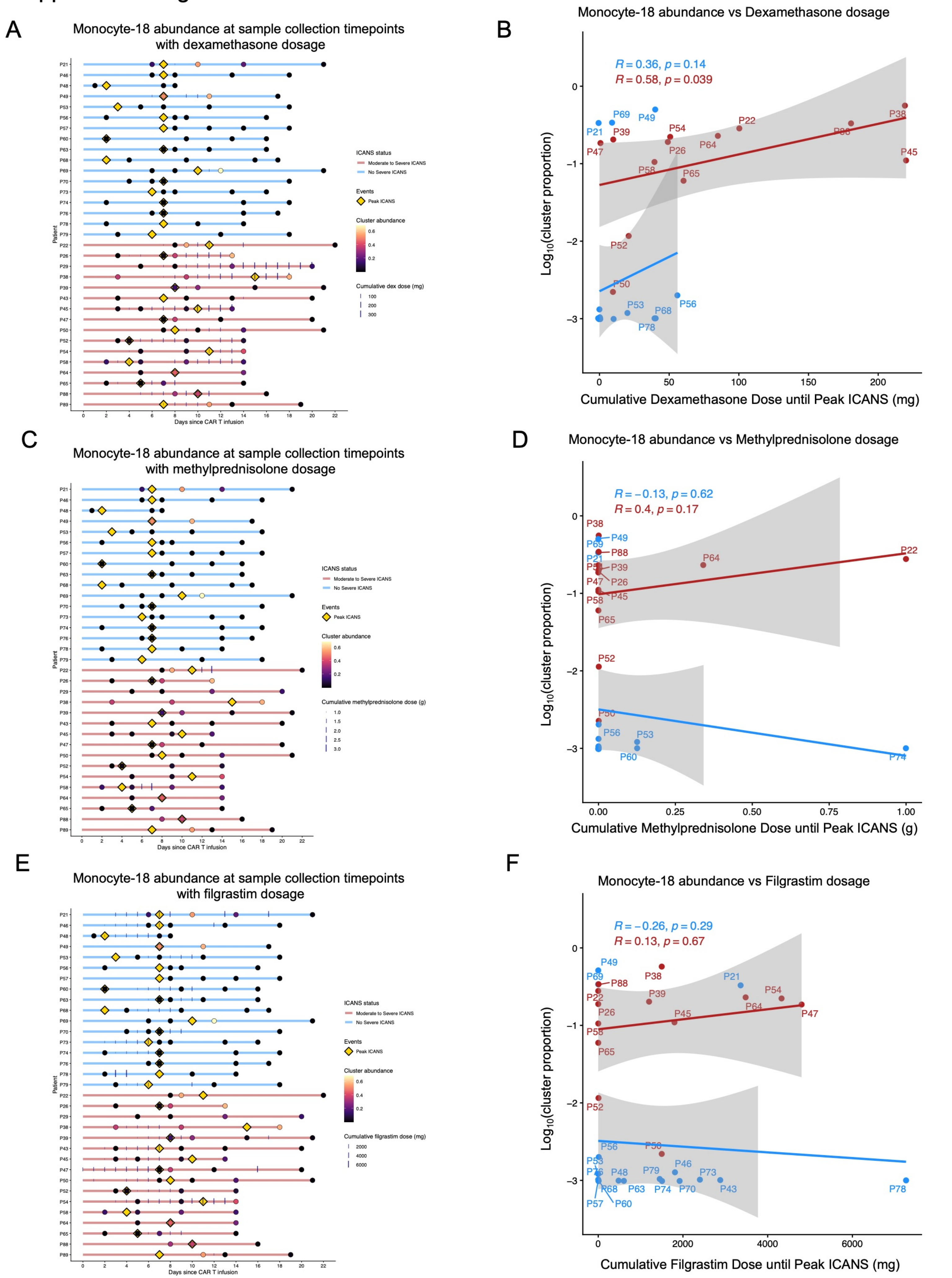

**Supplemental Figure 5. Association of corticosteroid and G-CSF exposure with CD163$^{+}$ Monocyte-18 abundance at peak ICANS.**

A. Swimmer plot depicting longitudinal Monocyte-18 abundance following CD19 CAR T-cell infusion in relation to dexamethasone administration. Horizontal lines represent the observation period for each patient from infusion through the final sample collection. Circles indicate sample collection timepoints and are scaled according to Monocyte-18 abundance. Yellow diamonds denote peak ICANS onset. Patients with non-severe ICANS are shown in blue and patients with moderate-to-severe ICANS are shown in red.
B. Relationship between cumulative dexamethasone exposure administered from CAR T-cell infusion through peak ICANS and Monocyte-18 abundance measured at peak ICANS. Each point represents an individual patient. Spearman correlation coefficients and corresponding P values are shown.
C. Swimmer plot depicting longitudinal Monocyte-18 abundance following CD19 CAR T-cell infusion in relation to methylprednisolone administration. Plot features are as described in (A).
D. Relationship between cumulative methylprednisolone exposure administered from CAR T-cell infusion through peak ICANS and Monocyte-18 abundance measured at peak ICANS. Each point represents an individual patient. Spearman correlation coefficients and corresponding P values are shown.
E. Swimmer plot depicting longitudinal Monocyte-18 abundance following CD19 CAR T-cell infusion in relation to filgrastim administration. Plot features are as described in (A).
F. Relationship between cumulative filgrastim exposure administered from CAR T-cell infusion through peak ICANS and Monocyte-18 abundance measured at peak ICANS. Each point represents an individual patient. Spearman correlation coefficients and corresponding P values are shown.

Supplemental Figure 6

A

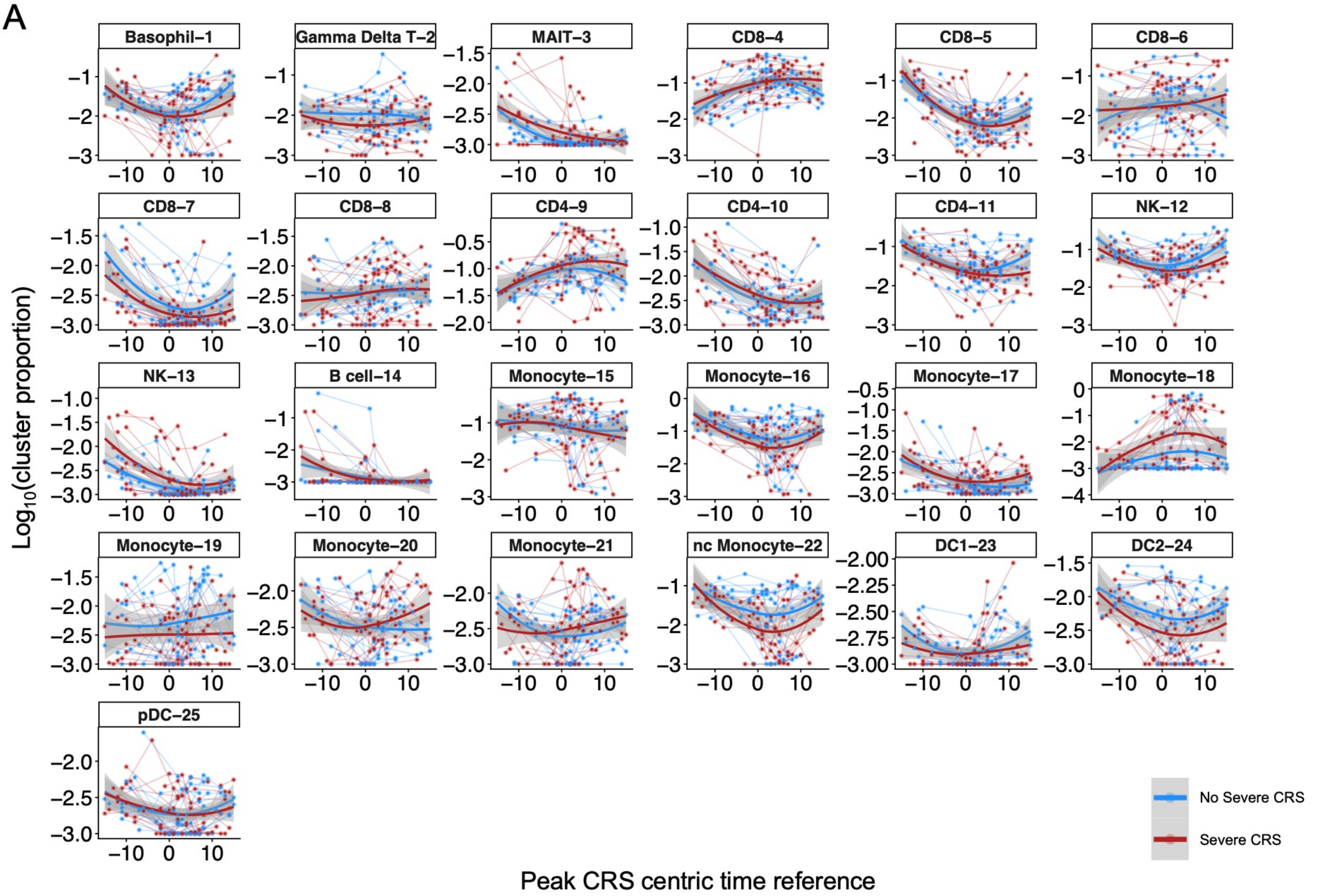


B

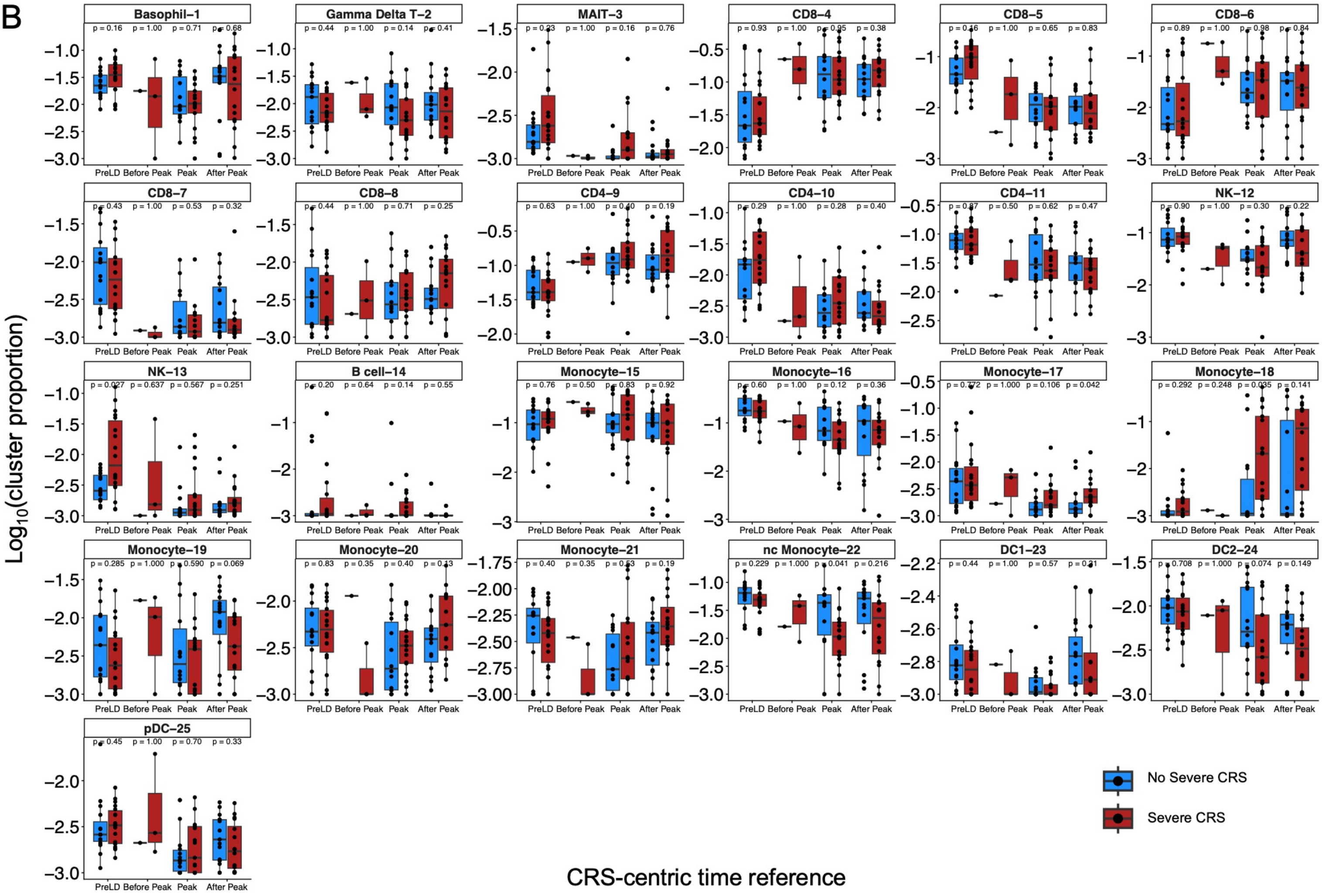

**Supplemental Figure 6. Longitudinal dynamics of mononuclear populations relative to peak CRS.**

A. Longitudinal trajectories of mononuclear population abundances identified by mass cytometry and plotted relative to patient-specific peak CRS onset. Individual points represent the frequency of each population within the $CD66b^{-}$ mononuclear compartment from samples collected throughout CD19 CAR T-cell therapy. Patients with non-severe CRS are shown in blue and patients with severe CRS are shown in red. Solid lines represent locally estimated scatterplot smoothing (LOESS) trends with 95% confidence intervals. Population labels correspond to unsupervised cluster identities generated from high-dimensional mass cytometry analysis.
B. Relative abundance of mononuclear populations across CRS-centric time windows, including pre-lymphodepletion (PreLD), Before Peak CRS, Peak CRS, and After Peak CRS, stratified by CRS severity. Patients with non-severe CRS are shown in blue and patients with severe CRS are shown in red. Each point represents an individual patient sample. Statistical comparisons were performed using two-sided Wilcoxon rank-sum tests, with P values indicated.

### Supplemental Figure 7

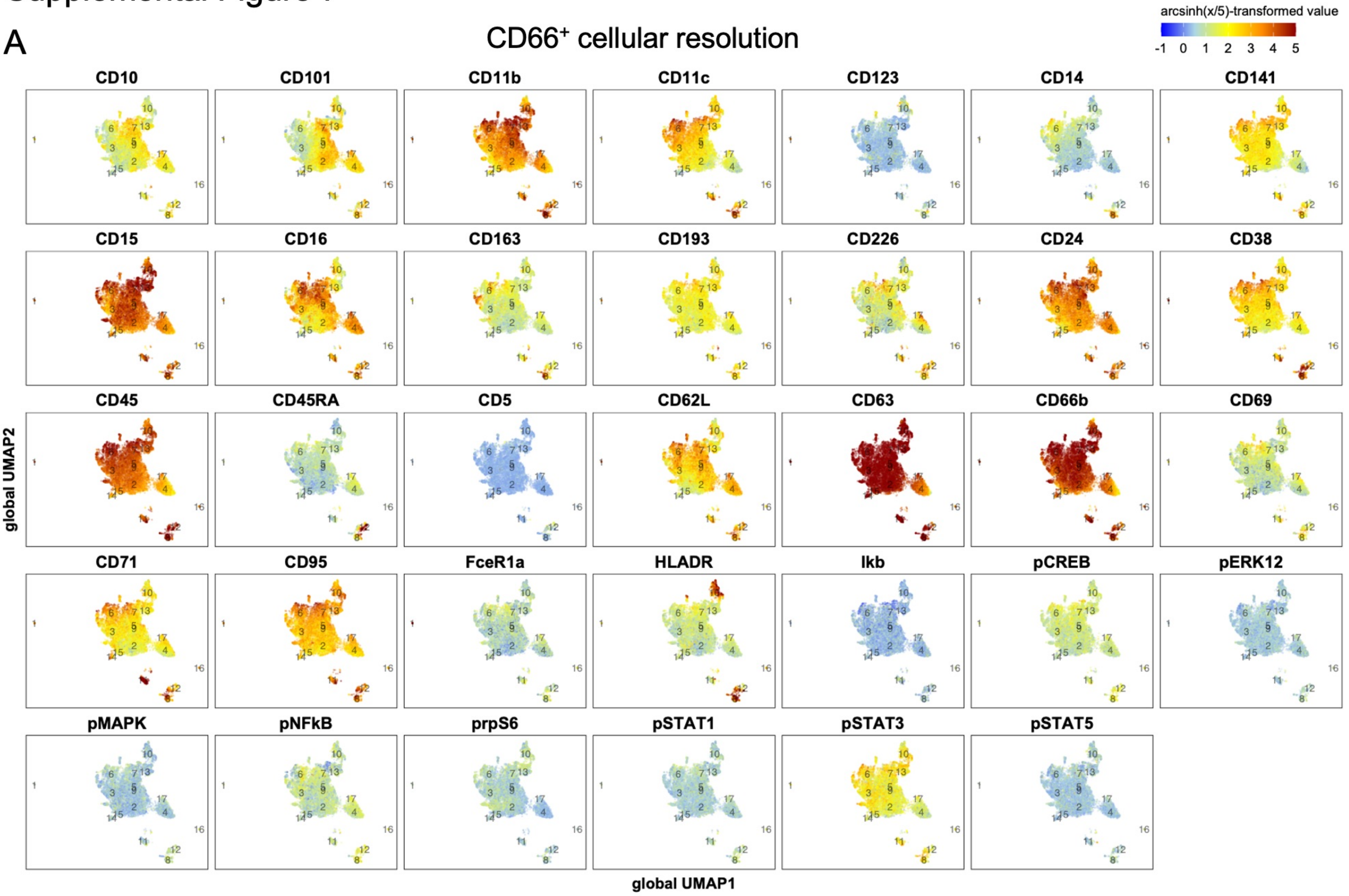


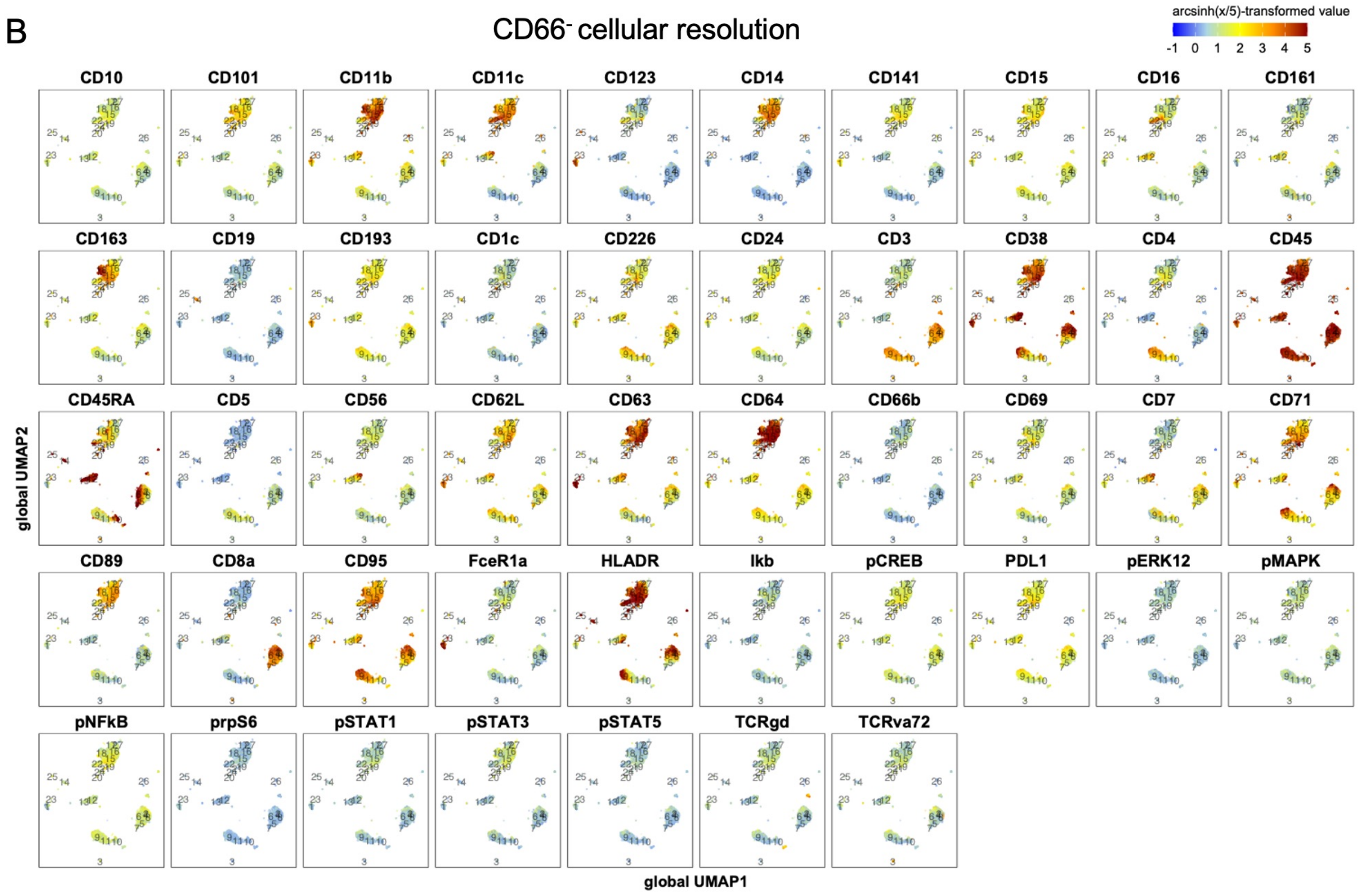

**Supplemental Figure 7. Phenotypic characterization of $CD66^{+}$ and $CD66b^{-}$ cell populations by mass cytometry marker expression**

A. UMAP feature plots of $CD66b^{+}$ granulocyte populations colored by arcsinh-transformed marker expression. Expression patterns highlight phenotypic heterogeneity across granulocyte subsets, including markers associated with maturation, activation, migration, and intracellular signaling.
B. UMAP feature plots of $CD66b^{-}$ mononuclear populations colored by arcsinh-transformed marker expression. Expression patterns define major immune populations, including T cells, B cells, monocytes, dendritic cells, and natural killer cells, as well as distinct activation and signaling states.

### Supplemental Figure 8

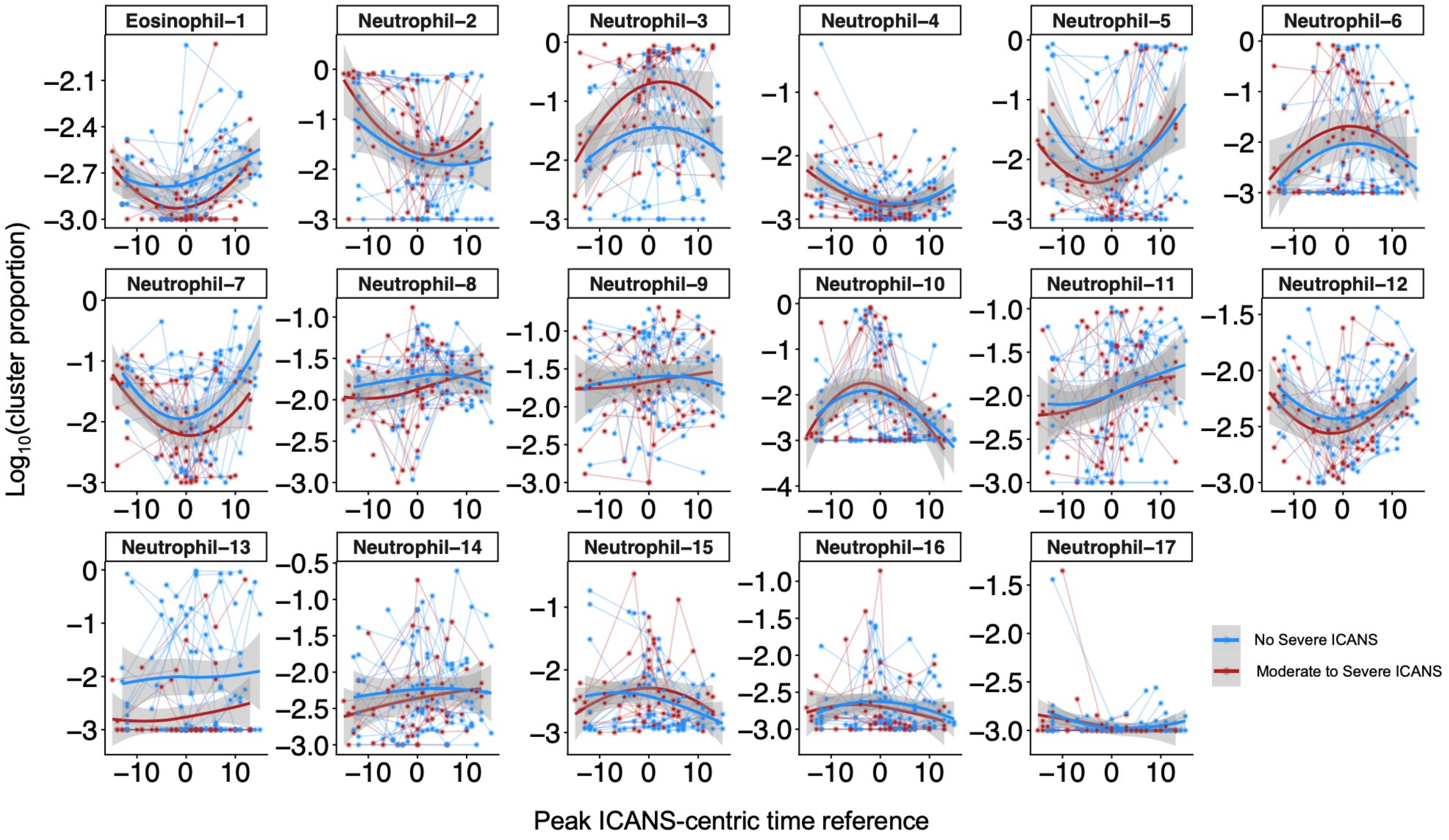

**Supplemental Figure 8. Longitudinal dynamics of granulocyte populations relative to peak ICANS onset.**

Longitudinal trajectories of granulocyte population abundances identified by mass cytometry and plotted relative to patient-specific peak ICANS onset. Individual points represent the frequency of each population within the $CD66b^{+}$ granulocyte compartment from samples collected throughout CD19 CAR T-cell therapy. Patients with non-severe ICANS are shown in blue and patients with moderate-to-severe ICANS are shown in red. Solid lines represent locally estimated scatterplot smoothing (LOESS) trends with 95% confidence intervals. Population labels correspond to unsupervised granulocyte cluster identities generated from high-dimensional mass cytometry analysis.

### Supplemental Figure 9

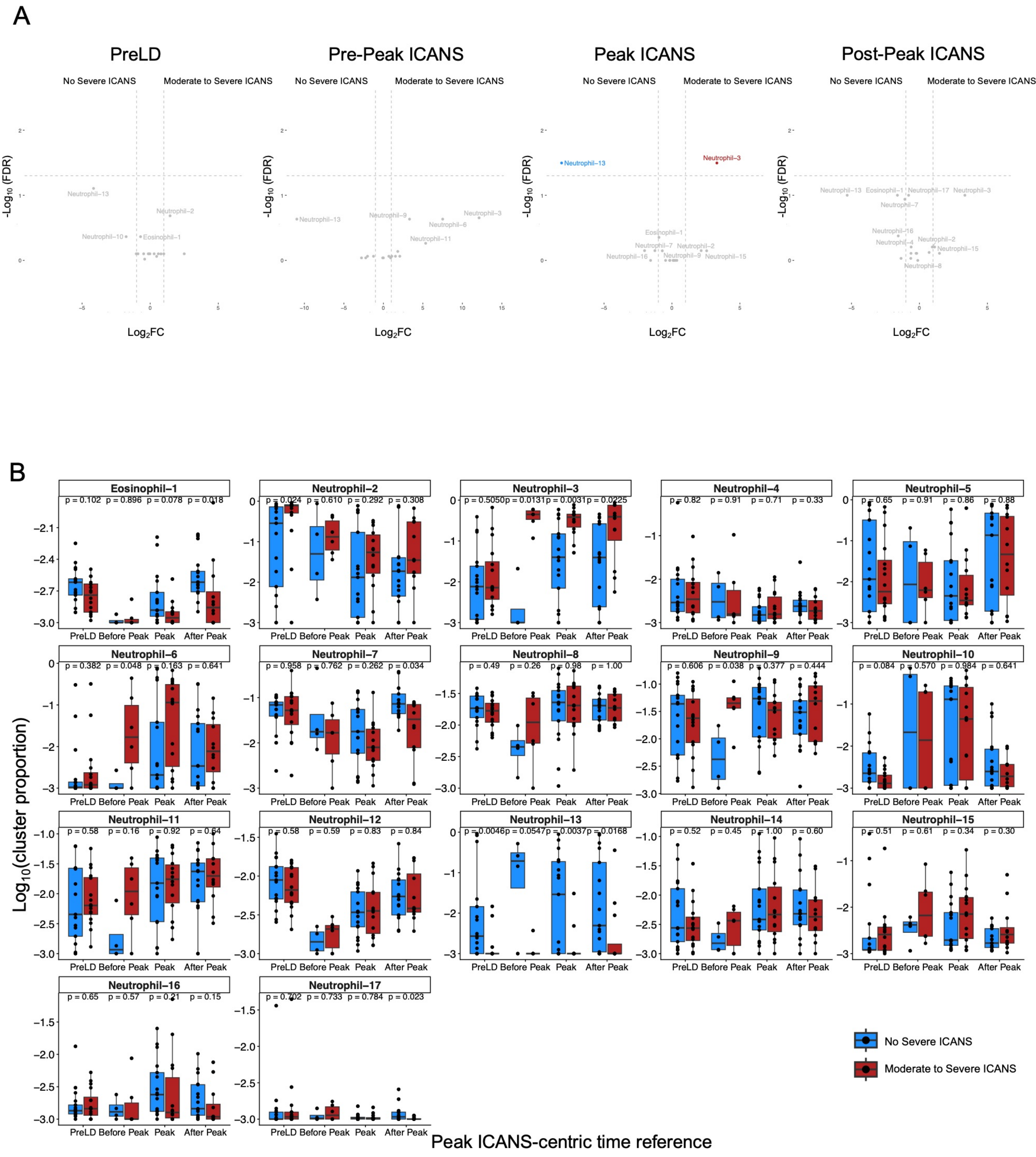

**Supplemental Figure 9. Differential abundance of granulocyte populations associated with ICANS severity.**

A. Volcano plots showing log2 fold changes in granulocyte population abundance between patients with moderate-to-severe ICANS and patients with non-severe ICANS at peak ICANS. Positive values indicate enrichment in moderate-to-severe ICANS, whereas negative values indicate enrichment in non-severe ICANS. Selected populations are annotated, and gray points represent populations that did not reach statistical significance.
B. Relative abundance of granulocyte populations across ICANS-centric time windows, including pre-lymphodepletion (PreLD), Before Peak ICANS, Peak ICANS, and After Peak ICANS, stratified by ICANS severity. Patients with non-severe ICANS are shown in blue and patients with moderate-to-severe ICANS are shown in red. Each point represents an individual patient sample. Statistical comparisons were performed using two-sided Wilcoxon rank-sum tests, with P values indicated.

### Supplemental Figure 10

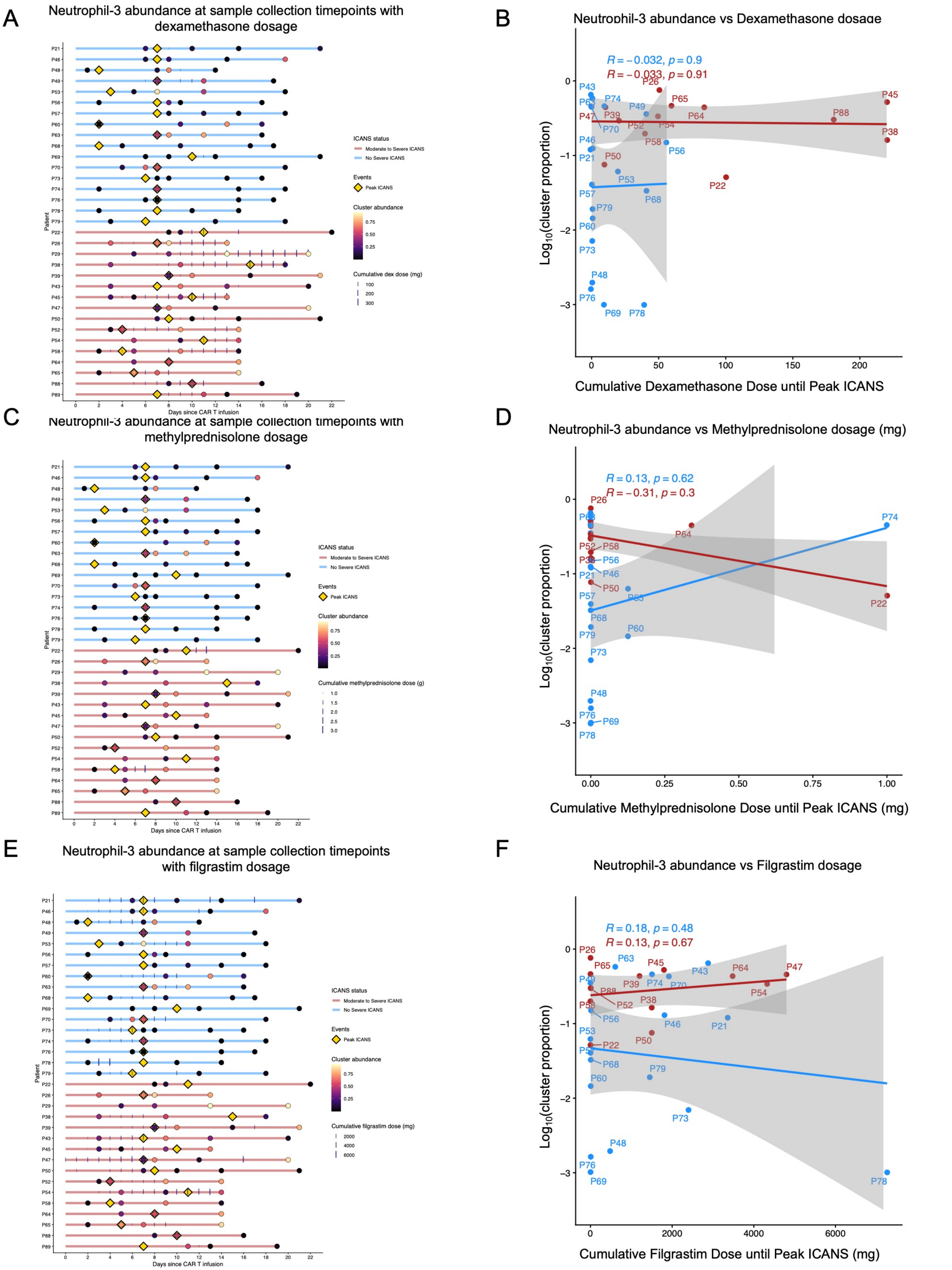

**Supplemental Figure 10. Association of corticosteroid and G-CSF exposure with CD10$^{-}$ Neutrophil-3 abundance at peak ICANS.**

A. Swimmer plot depicting longitudinal Neutrophil-3 abundance following CD19 CAR T-cell infusion in relation to dexamethasone administration. Horizontal lines represent the observation period for each patient from infusion through the final sample collection. Circles indicate sample collection timepoints and are scaled according to Neutrophil-3 abundance. Yellow diamonds denote peak ICANS onset. Patients with non-severe ICANS are shown in blue and patients with moderate-to-severe ICANS are shown in red.
B. Relationship between cumulative dexamethasone exposure administered from CAR T-cell infusion through peak ICANS and Neutrophil-3 abundance measured at peak ICANS. Each point represents an individual patient. Spearman correlation coefficients and corresponding P values are shown.
C. Swimmer plot depicting longitudinal Neutrophil-3 abundance following CD19 CAR T-cell infusion in relation to methylprednisolone administration. Plot features are as described in (A).
D. Relationship between cumulative methylprednisolone exposure administered from CAR T-cell infusion through peak ICANS and Neutrophil-3 abundance measured at peak ICANS. Each point represents an individual patient. Spearman correlation coefficients and corresponding P values are shown.
E. Swimmer plot depicting longitudinal Neutrophil-3 abundance following CD19 CAR T-cell infusion in relation to filgrastim administration. Plot features are as described in (A).
F. Relationship between cumulative filgrastim exposure administered from CAR T-cell infusion through peak ICANS and Neutrophil-3 abundance measured at peak ICANS. Each point represents an individual patient. Spearman correlation coefficients and corresponding P values are shown.

Supplemental Figure 11

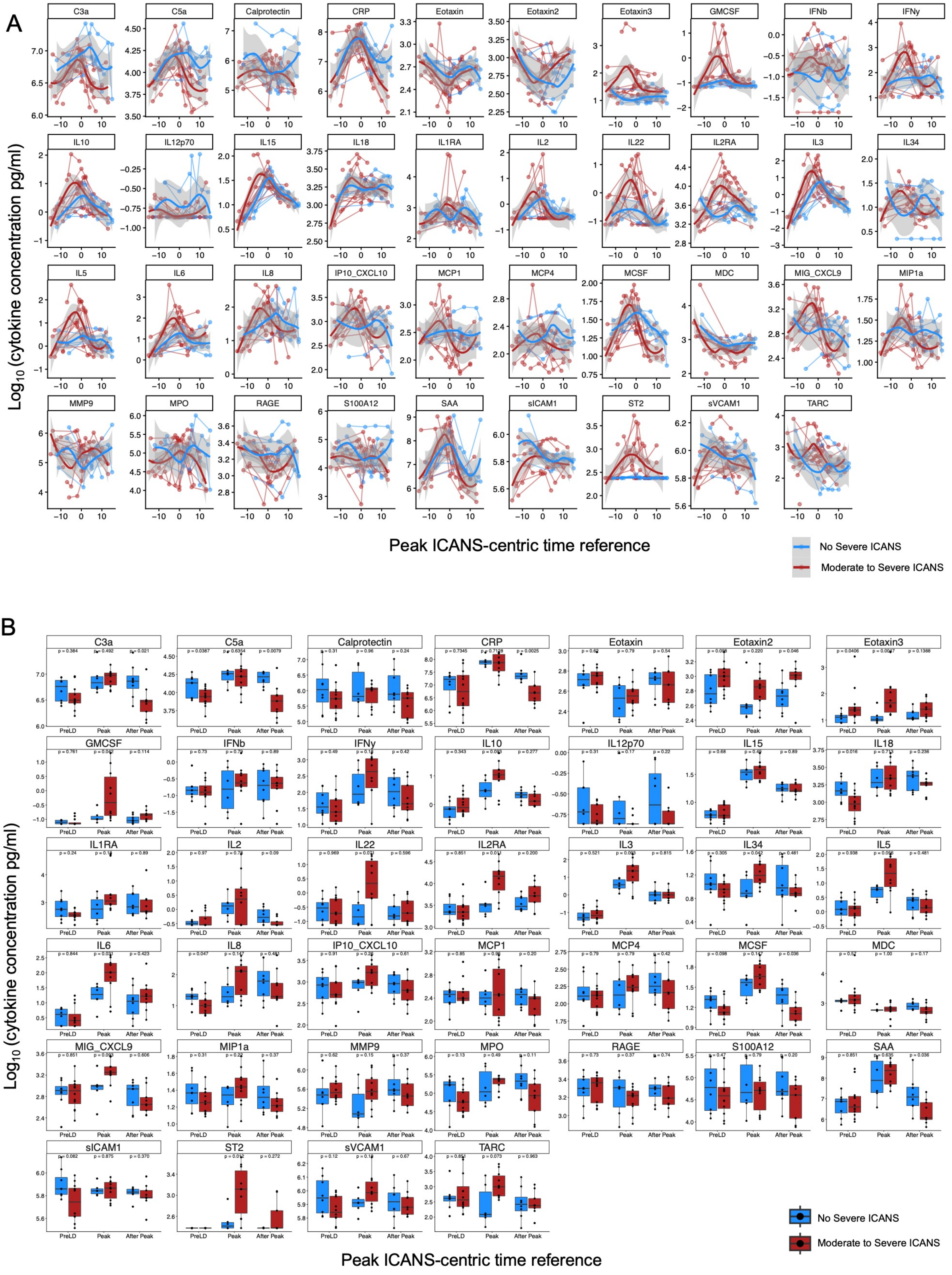

**Supplemental Figure 11. Longitudinal dynamics of circulating serum proteins relative to peak ICANS onset.**

A. Longitudinal trajectories of serum protein concentrations plotted relative to patient-specific peak ICANS onset. Individual points represent serum samples collected throughout CD19 CAR T-cell therapy. Patients with non-severe ICANS are shown in blue and patients with moderate-to-severe ICANS are shown in red. Solid lines represent locally estimated scatterplot smoothing (LOESS) trends with 95% confidence intervals.
B. Serum protein concentrations across ICANS-centric time windows, including pre-lymphodepletion (PreLD), Peak ICANS, and After Peak ICANS, stratified by ICANS severity. Patients with non-severe ICANS are shown in blue and patients with moderate-to-severe ICANS are shown in red. Each point represents an individual patient sample. Statistical comparisons were performed using two-sided Wilcoxon rank-sum tests, with P values indicated.

Supplemental Figure 12

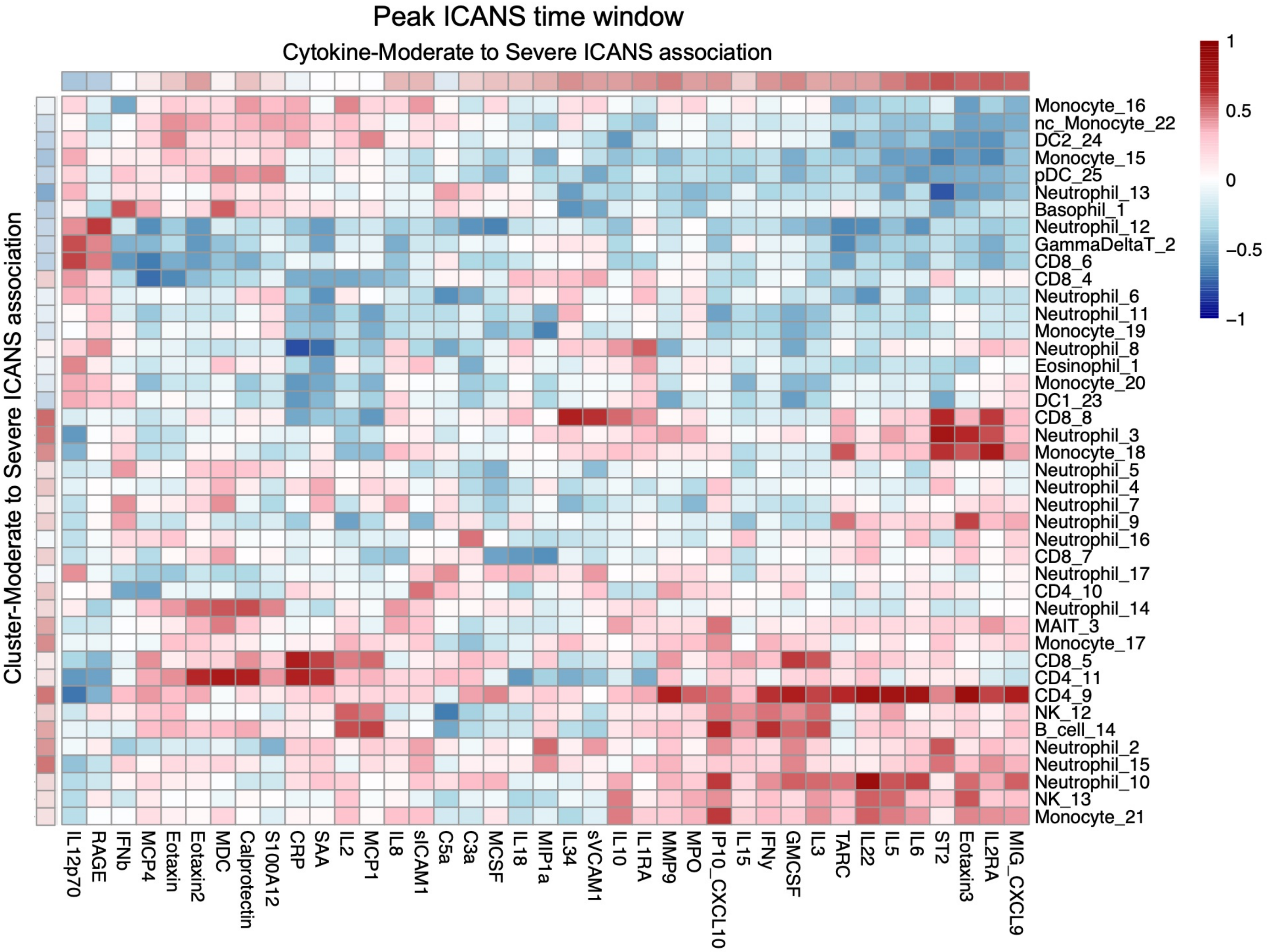

**Supplemental Figure 12. Cellular and proteomic correlations in the context of ICANS at peak toxicity**

Correlation matrix depicting associations between serum cytokines, soluble mediators, and protein concentrations with mononuclear and granulocyte cluster abundances at peak ICANS. Rows represent soluble proteins and columns represent mononuclear and granulocyte populations. Red indicates positive correlations, whereas blue indicates negative correlations. Color intensity reflects the strength of the correlation between cellular population abundance and soluble mediator concentration at peak neurotoxicity as measured by Spearman correlation coefficient (ρ).

### Supplemental Figure 13

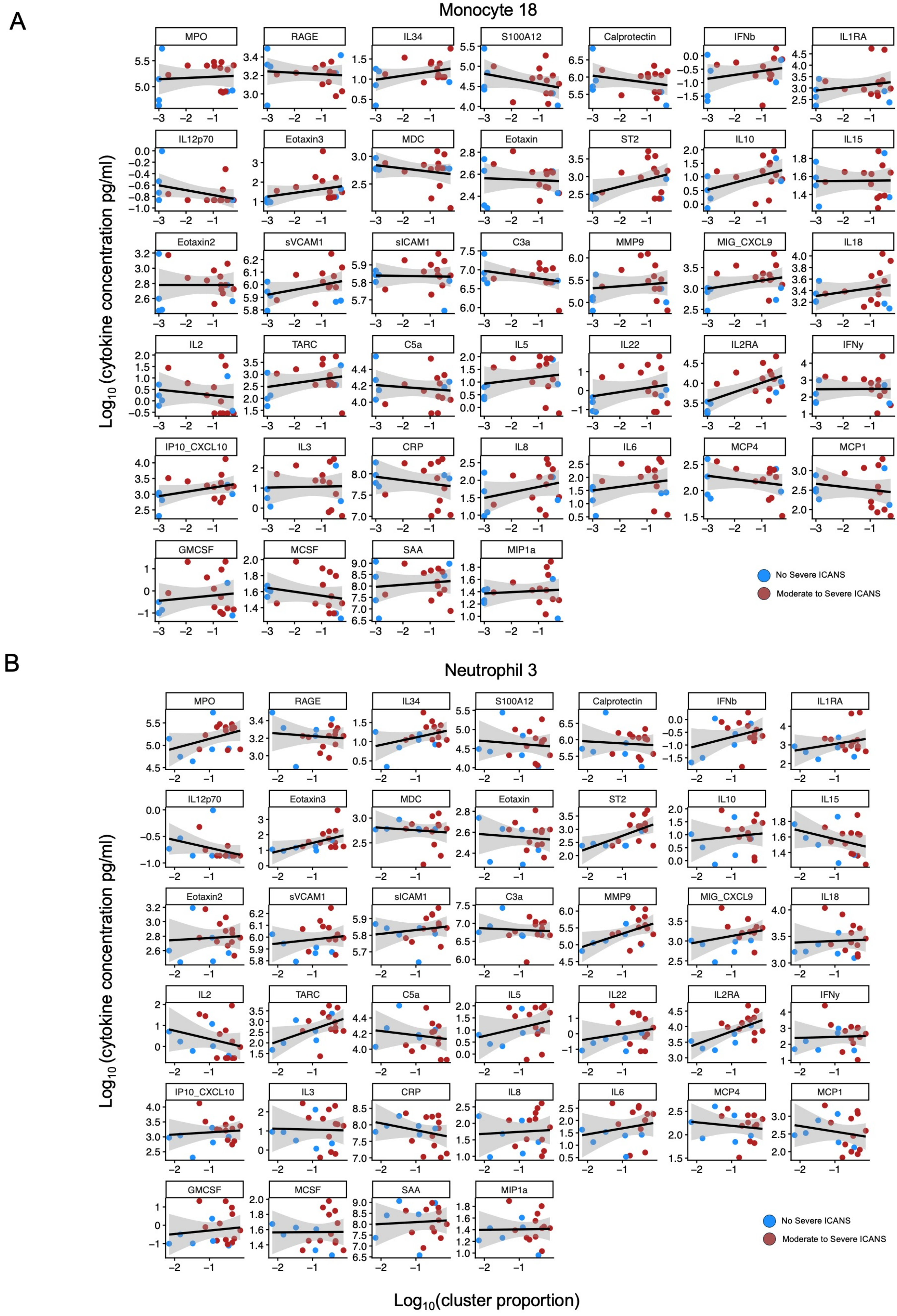

**Supplemental Figure 13. Monocyte-18 and Neutrophil-3 cytokine correlations at peak ICANS**

A. Scatter plots depicting correlations between Monocyte-18 abundance and serum cytokine, chemokine, and soluble mediator concentrations measured at peak ICANS. Each point represents an individual patient sample collected within the peak neurotoxicity window. Patients are stratified by ICANS severity, with moderate to severe ICANS shown in red and no severe ICANS shown in blue.
B. Scatter plots depicting correlations between Neutrophil-3 abundance and serum cytokine, chemokine, and soluble mediator concentrations measured at peak ICANS. Each point represents an individual patient sample collected within the peak neurotoxicity window. Patients are stratified by ICANS severity, with moderate to severe ICANS shown in red and no severe ICANS shown in blue.

Supplemental Figure 14

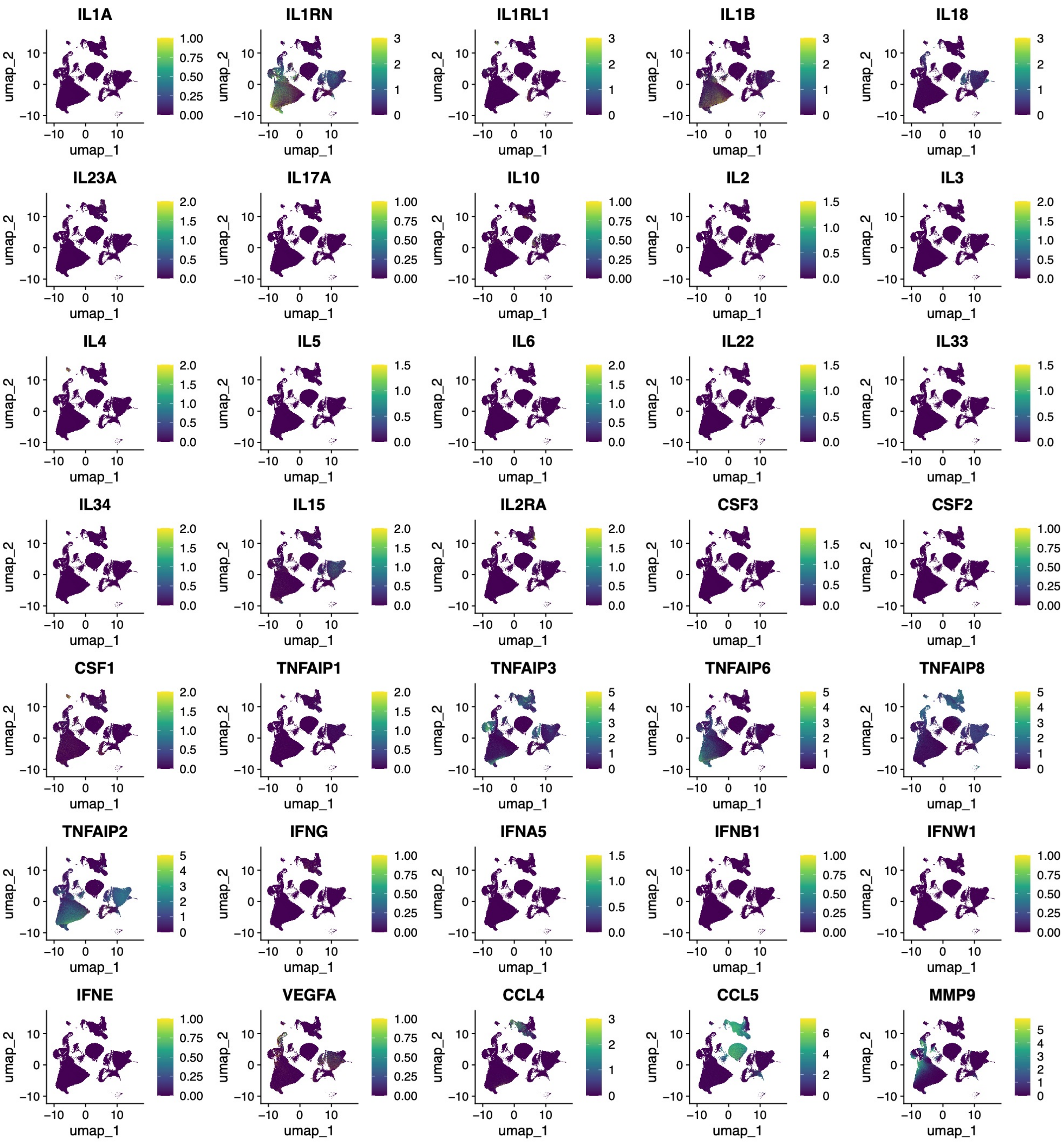

**Supplemental Figure 14. Cytokine gene expression across immune populations in the validation cohort.**

Feature plots showing the expression of selected cytokine genes across immune populations identified by 10x Genomics GEM-X FLEX APEX profiling. Cells are colored according to normalized gene expression, highlighting the cellular distribution of cytokine transcripts across major immune subsets.

Supplemental Figure 15

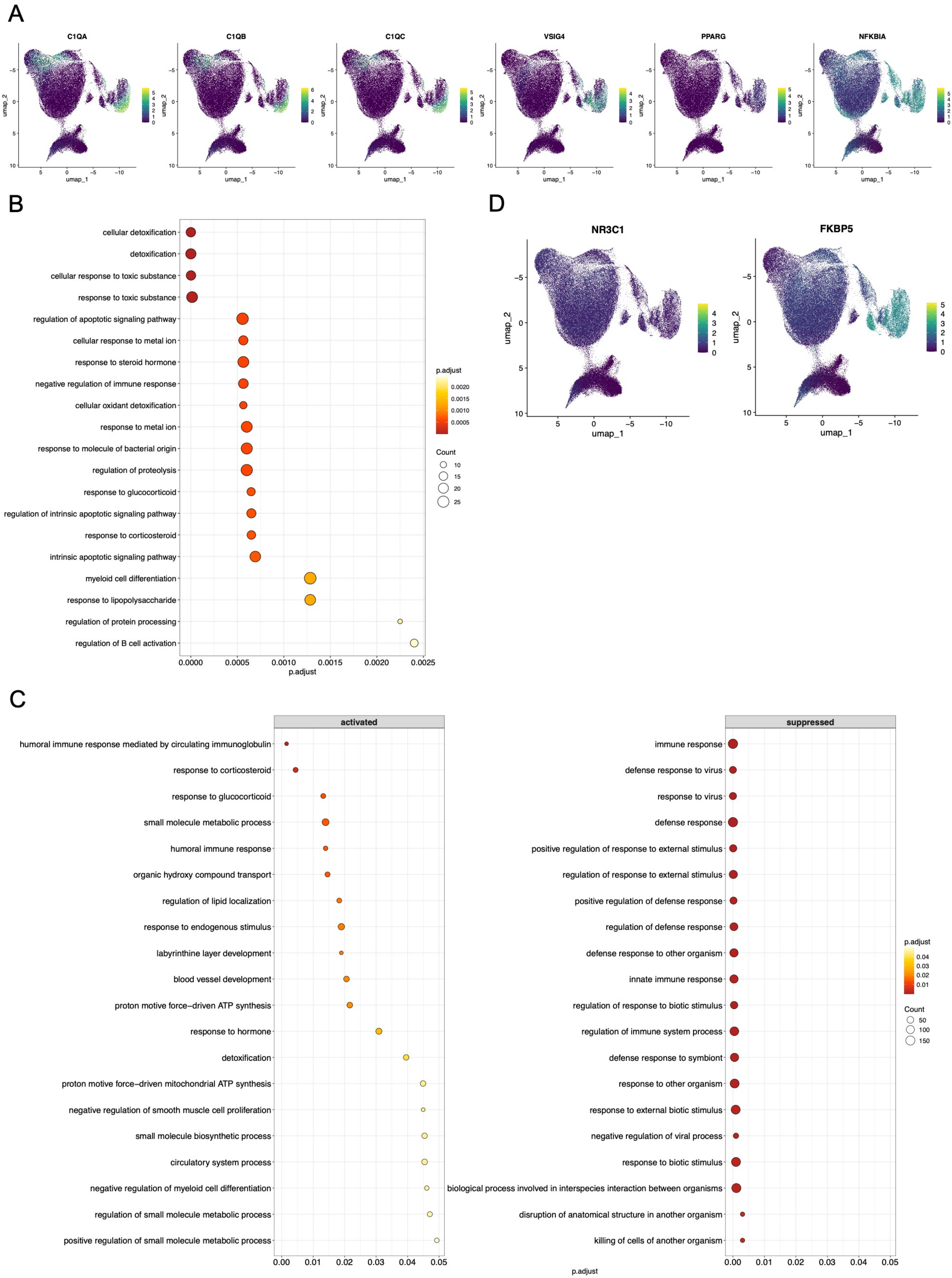

**Supplemental Figure 15. Monocyte-3 exhibits an immunoregulatory and glucocorticoid-responsive transcriptional profile**

A. Feature plots showing expression of representative genes enriched in Monocyte-3, including *C1QA, C1QB, C1QC, VSIG4, PPARG,* and *NFKBIA*
B. Gene Ontology enrichment analysis of genes overexpressed in Monocyte-3, demonstrating enrichment of pathways associated with detoxification, cellular detoxification, and responses to toxic substances.
C. Activated and suppressed biological pathways identified by gene set enrichment analysis of Monocyte-3 relative to other monocyte populations. Activated pathways include corticosteroid and glucocorticoid response programs, whereas suppressed pathways are associated with immune and antiviral responses.
D. Feature plots showing expression of NR3C1 and the glucocorticoid-responsive gene FKBP5 across monocyte populations, demonstrating preferential enrichment of FKBP5 within Monocyte-3.

#### Supplemental Figure 16

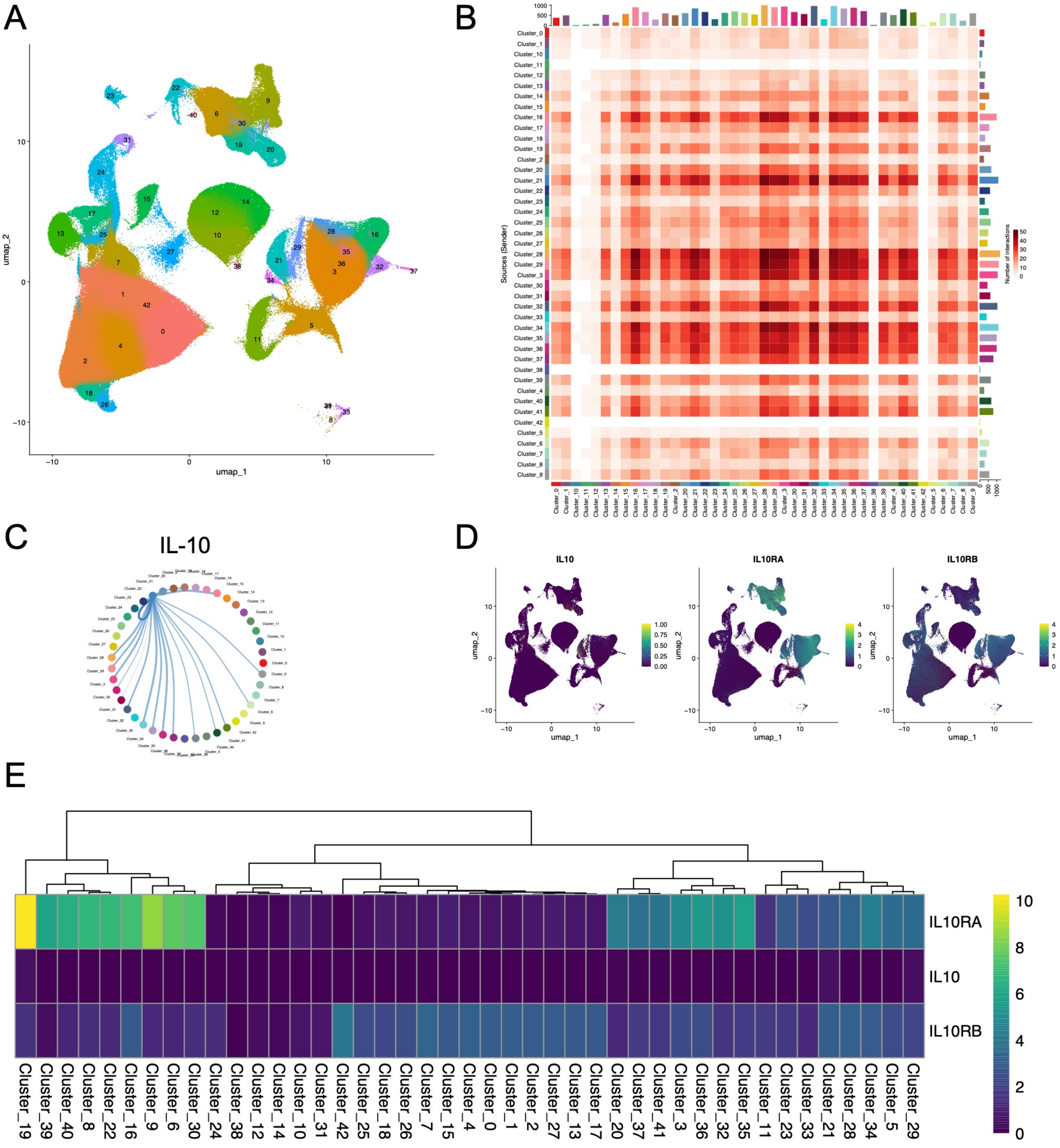

**Supplemental Figure 16. CD163$^{+}$ monocytes are the predominant source of IL10 transcripts and predicted IL-10 signaling.**

A. UMAP visualization of immune populations identified by 10x Genomics GEM-X FLEX APEX profiling of fixed whole blood samples.
B. Heatmap showing inferred cell–cell communication networks generated using CellChat analysis. Rows represent signaling source (sender) populations and columns represent signaling target (receiver) populations. Color intensity reflects the number of predicted ligand–receptor interactions between cell populations.
C. Circle plot depicting predicted IL-10 signaling interactions among immune populations based on CellChat communication probabilities.
D. Feature plots characterizing expression of *IL10, IL10RA* and *IL10RB* across all cells
E. Heatmap displaying scaled expression of *IL10*, *IL10RA*, and *IL10RB* across resolved clusters.

Supplemental Figure 17

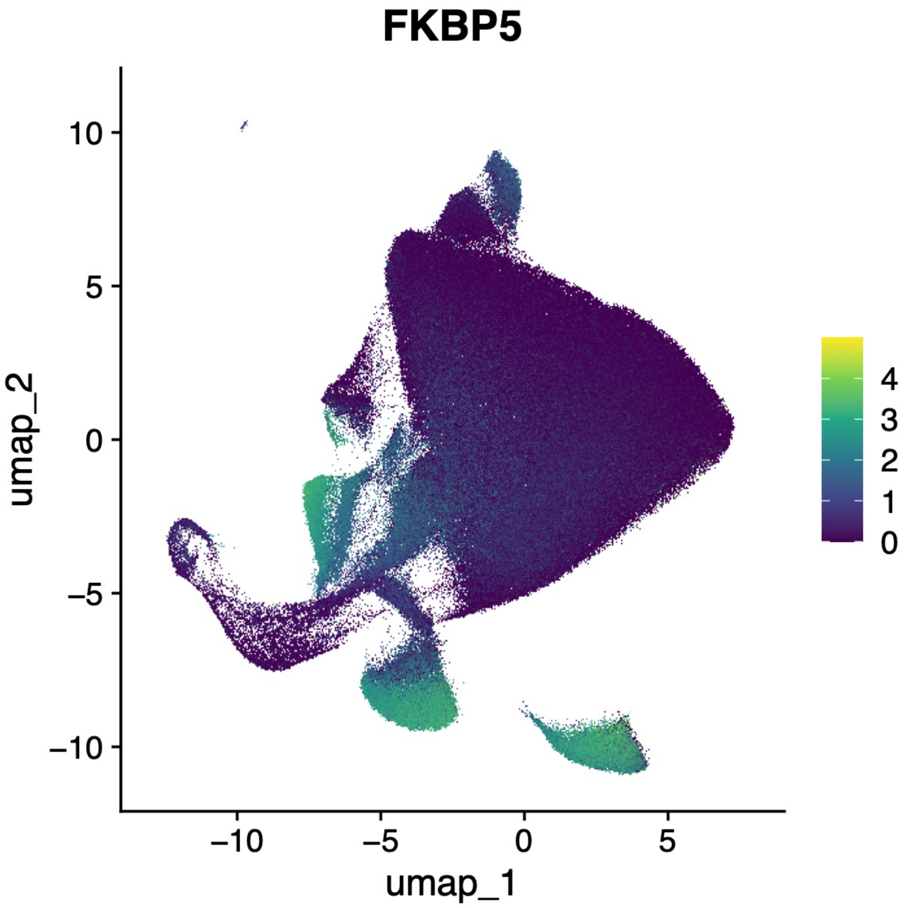

**Supplementary Figure 17. FKBP5 expression is enriched in granulocyte populations associated with moderate-to-severe ICANS.**

Feature plot showing FKBP5 expression across granulocyte populations identified by 10x Genomics FLEX APEX profiling.
