## Supplemental Tables for "Coordinated expansion of CD163⁺ monocytes and immature CD177⁺ neutrophils marks severe neurotoxicity after CD19 CAR T cell therapy"

Supplemental Table 1

| Characteristic | Discovery Cohort N = 33[1] | Validation Cohort N = 12[1] |
|---|---|---|
| **Age at CAR T-cell infusion (years)** | | |
| Median (Q1, Q3) | 67.1 (57.6, 71.5) | 71.7 (68.0, 76.8) |
| Min - Max | 33.5 - 84.3 | 64.5 - 81.9 |
| **Sex** | | |
| Female | 14 (42%) | 6 (50%) |
| Male | 19 (58%) | 6 (50%) |
| **Diagnosis** | | |
| Acute lymphoblastic leukemia | 1 (3%) | 0 (0%) |
| Follicular lymphoma | 1 (3%) | 2 (17%) |
| Large B-cell lymphoma | 29 (88%) | 5 (42%) |
| Mantle cell lymphoma | 2 (6%) | 5 (42%) |
| **CAR T-cell product** | | |
| Axicabtagene ciloleucel | 18 (55%) | 0 (0%) |
| Brexucabtagene autoleucel | 3 (9%) | 0 (0%) |
| Lisocabtagene maraleucel | 6 (18%) | 12 (100%) |
| Tisagenlecleucel | 6 (18%) | 0 (0%) |

[1] n (%)

Supplemental Table 2

| Target | Clone | Metal Isotope | Source | Part number |
|---|---|---|---|---|
| CD45 | HI3O | 89 Y | Standard Biotools | 3089003 |
| CD4 | RPA-T4 | 110 Cd | BioLegend | 300502 |
| CD8a | RPA-T8 | 111 Cd | BioLegend | 301002 |
| CD7 | CD7-6B7 | 112 Cd | BioLegend | 343102 |
| CD63 | H5C6 | 113 Cd | BD Pharmingen | 556019 |
| BDCA3/CD141 | 1A4 | 114 Cd | BD Pharmingen | 230282 |
| CD66b | G10F5 | 116 Cd | BioLegend | 305102 |
| CD10 | HI10a | 141 Pr | BioLegend | 312202 |
| CD101 | BB27 | 142 Nd | ThermoFisher | 14-1019-82 |
| CD123 | 6H6 | 143 Nd | Standard Biotools | 3143014B |
| TCRgdAnti-FITC | 5A6.E9/FIT-22 | 144 Nd | ThermoFischer/Standard Biotools | MHGD01/3144006B |
| CD163 | GHI/61 | 145 Nd | Standard Biotools | 3145010B |
| CD69 | FN50 | 146 Nd | BioLegend | 310902 |
| CD16 | 3G8 | 148 Nd | Standard Biotools | 3148004B |
| CD24 | ML5 | 149 Sm | BioLegend | 311102 |
| FcεR1α | AER-37 | 150 Sm | Standard Biotools | 3150027B |
| CD226 | DNAM-1 | 151 Eu | BioLegend | 338302 |
| CD62L | DREG-56 | 152 Sm | BioLegend | 304802 |
| CD38 | HIT2 | 154 Sm | BioLegend | 303502 |
| CD95 | DX2 | 155 Gd | BioLegend | 305602 |
| TCRva7.2 | 3C10 | 156 Gd | BioLegend | 351702 |
| CD15 | W6D3 | 157 Gd | BioLegend | 323002 |
| CD235AB | HIR2 | 160 Dy | BioLegend | 306602 |
| CD161 | HP-3G10 | 161 Dy | BioLegend | 339902 |
| CD71 | OKT-9 | 162 Dy | ThermoFisher | 14-0719-82 |
| CD1c | L161 | 163 Dy | BioLegend | 331502 |
| CD64 | 10.1 | 165 Ho | BioLegend | 305002 |
| CD19 | HIB19 | 168 Er | BioLegend | 302202 |
| CD45RA | HI100 | 169 Tm | Standard Biotools | 3169008B |
| PDL1/CD274 | 29E.2A3 | 170 Er | BioLegend | 329702 |
| CD56 | HCD56 | 171 Yb | BioLegend | 318345 |
| CD193 | 5E8 | 172 Yb | BioLegend | 310702 |
| HLA-DR | L243 | 173 Yb | Standard Biotools | 3173005B |
| CD89 | A59 | 174 Yb | Standard Biotools | 3174012B |
| CD3 | UCHT1 | 194 Pt | BioLegend | 300402 |
| CD5 | UCHT2 | 195 Pt | BioLegend | 300602 |
| CD14 | HCD14 | 198 Pt | BioLegend | 325602 |
| CD11b | ICRF44 | 209 Bi | Standard Biotools | 3209003B |
| CD11c | 3.9 | 106 Cd | BioLegend | 301602 |
| pSTAT1 | 58D6 | 153 Eu | Standard Biotools | 3153003A |
| pSTAT3 | 4/P-STAT3 | 158 Gd | Standard Biotools | 3158005A |
| pSTAT5 | 47 | 147 Sm | Standard Biotools | 3147012A |
| pCREB | 87G3 | 176 Lu | Standard Biotools | 3176005A |
| prpS6 | N7-548 | 175 Lu | Standard Biotools | 3175009A |
| pMAPKAPK2 | 27B7 | 159 Tb | Standard Biotools | 3159010A |
| pNFκB | K10x | 166 Er | Standard Biotools | 3166006A |
| pERK1/2 | D1314.4E | 167 Er | Standard Biotools | 3167005A |
| Ikb | L35A5 | 164 Dy | Standard Biotools | 3164004A |

Supplemental Table 3

| Assay | Vendor | CATALOG # | Format | Analyte | Sample dilution |
|---|---|---|---|---|---|
| A | MSD | K15067M-2 | UPLEX Custom Biomarker | GM-CSF, IL-10, MIP1a, IL-2, IL-22, IL-12p70, IL-5, MCP4, IL-4, IL-13 | 1 |
| B | MSD | K15067M-2 | UPLEX Custom Biomarker | IFNy, IL-18, IL-1RA, IL-15, IL-6, IL-8, IP-10, MCP1, sIL-2Ra, MIG | 4 |
| C | MSD | K151198D-2 | V-PLEX Vascular Injury | CRP, SAA, ICAM-1, VCAM-1 | 10,000 |
| D | MSD | K151P3S-2 | SPLEX | IFN-a2a | 1 |
| E | MSD | K151ADQS-2 | SPLEX | IL-3 | 1 |
| F | MSD | K151ADRS-2 | SPLEX | IFNb | 2 |
| G | R&D | DY119 | ELISA | IL-18BP | 40 or 200 |
| H | MSD | K151AEM-2 | UPLEX Custom Immuno-oncology | MMP9(total), S100A12 | 100 |
| I | MSD | K1514ER-2 | RPLEX | MPO | 20 |
