## Supplemental Table Legends for "Coordinated expansion of CD163⁺ monocytes and immature CD177⁺ neutrophils marks severe neurotoxicity after CD19 CAR T cell therapy"

**Supplemental Table 1. Clinical characteristics of the discovery and validation cohorts.**

Demographic and disease characteristics of patients included in the CyTOF discovery cohort (n = 33) and the 10x Genomics FLEX APEX validation cohort (n = 12). Age at CAR T-cell infusion is presented as median with interquartile range (Q1–Q3) and minimum–maximum values. Categorical variables are presented as number of patients (%). CAR T-cell products included axicabtagene ciloleucel, brexucabtagene autoleucel, lisocabtagene maraleucel, and tisagenlecleucel.

**Supplemental Table 2. Mass cytometry antibody panel used for longitudinal immune profiling following CD19 CAR T cell therapy**

Antibodies targeting lineage, activation, adhesion, checkpoint, cytokine signaling, and intracellular phosphoprotein markers were used for high-dimensional mass cytometry analysis of fixed whole blood samples collected longitudinally from patients undergoing CD19 CAR T cell therapy. The panel included markers for monocytes, granulocytes, dendritic cells, T cells, B cells, NK cells, and intracellular signaling pathways relevant to immune activation and inflammation. Listed are the target antigen, antibody clone, metal isotope conjugate, reagent source, and catalog/part number for each antibody used in the study.

**Supplemental Table 3. Meso Scale Discovery (MSD) assay panels used for longitudinal serum proteomic profiling following CD19 CAR T cell therapy**

Serum proteins were quantified using custom and commercially available Meso Scale Discovery (MSD) multiplex immunoassay platforms. Assays included inflammatory cytokines, chemokines, vascular injury markers, immune activation markers.
